# Synthetic budding morphogenesis by optogenetic receptor tyrosine kinase signaling

**DOI:** 10.64898/2026.03.31.715459

**Authors:** Louis S. Prahl, Ronald Canlla, Aria Zheyuan Huang, Daniel S. Alber, Sandra L. Shefter, Sachin N. Davis, Samuel H. Grindel, Zikang Dennis Huang, Thomas R. Mumford, William Benman, Lukasz J. Bugaj, Kyle W. McCracken, Alex J. Hughes

**Affiliations:** Department of Bioengineering, University of Pennsylvania, Philadelphia, PA USA; Center for Soft & Living Matter, University of Pennsylvania, Philadelphia, PA USA; Bioengineering Graduate Group, University of Pennsylvania, Philadelphia, PA USA; Center for Engineering Mechanobiology, University of Pennsylvania, Philadelphia, PA USA; Center for Stem Cell and Organoid Medicine (CuSTOM), Cincinnati Children’s Hospital Medical Center, Cincinnati, OH USA; Divisions of Nephrology and Developmental Biology, Department of Pediatrics, Cincinnati Children’s Hospital Medical Center, Cincinnati, OH USA; Department of Pediatrics, University of Cincinnati College of Medicine, Cincinnati, OH USA; Department of Cell & Developmental Biology, University of Pennsylvania, Philadelphia, PA USA; Cell & Developmental Biology Graduate Group, Perelman School of Medicine, University of Pennsylvania, Philadelphia, PA USA; Institute for Regenerative Medicine, Perelman School of Medicine, University of Pennsylvania, Philadelphia, PA USA; Center for Precision Engineering for Health (CPE4H), University of Pennsylvania, Philadelphia, PA USA; Current address: BioFrontiers Institute and Department of Chemical and Biological Engineering, University of Colorado, Boulder, CO USA; Current address: Merge Labs, Brisbane, CA USA

**Keywords:** Epithelial cells, branching morphogenesis, synthetic morphogenesis, development, organogenesis, kidney, organoid, optogenetics, receptor tyrosine kinase signaling

## Abstract

The mammalian kidney relies on a branched network of collecting ducts for fluid transport and homeostasis. Replicating this network *in vitro* would parallelize function in synthetic replacement kidneys, yet current organoids have limited branching capacity. Here, we establish a developmentally-informed strategy to control organoid budding through optogenetic control of a receptor tyrosine kinase, RET. We first show pharmacological manipulation of RET signaling controls the extent of branching in mouse embryonic kidneys and human stem cell-derived kidney organoids. Next, we develop an optogenetic RET receptor (optoRET) that signals in a ligand-independent manner via blue light-mediated clustering. Epithelial cells expressing optoRET reproduce stereotyped RET signaling, scattering, and symmetry breaking in response to blue light. Human kidney organoids undergo budding with controllable orientation in response to spatially patterned optoRET stimulation. Our results establish ligand-free optogenetic control of branching and inspire new synthetic biology strategies for epithelial organoid design.

**Highlights:** GDNF-RET controls branching and tip cell state in mouse and human kidney tissues.

OptoRET reproduces endogenous RET signaling and morphogenesis in cell lines.

OptoRET enables ligand-free budding in human renal epithelial organoids.

Spatially patterned optoRET stimulation controls budding orientation.

## Introduction

Several epithelial organs including the lung, mammary gland, and kidney derive their function from thousands to millions of ducts and functional cellular units such as alveoli, acini, and nephrons ^1^. Branching morphogenesis builds and organizes these structures, allowing many parallel units to connect to a single exit point. One strategy to reconstitute this for tissue engineering is to harness native collective cell behaviors that drive morphogenesis ^2^.

Optogenetics is particularly suited to multicellular engineering ^3–5^ because light stimuli can be delivered or removed nearly instantaneously with subcellular spatial precision across centimeter-scale tissues to achieve ‘remote control’ of cell signaling and tissue morphogenesis^6^. Receptor tyrosine kinases (RTKs) are prime targets for optogenetic morphogenesis, given their *in vivo* roles in regulating spatially coordinated processes such as collective migration ^7^, epithelial branching ^8^, and positional control of morphogen signaling ^9^. Additionally, researchers have now functionally reconstituted the signaling dynamics of several endogenous pathways using optogenetic RTKs (optoRTKs) ^10–12^.

Recapitulating the tree-like architecture of branching epithelial ducts found in developing organs is a grand challenge for organoid engineering. Branching signals and kinematics are organ-specific but may share common motifs ^1^. One of these is a tip-localized epithelial progenitor cell population that drives branching through sustained RTK interactions with ligands in the surrounding mesenchyme ^13^. In the embryonic kidney, ureteric bud tip (and not trunk) cells express the <u>RE</u>arranged during <u>T</u>ransfection (RET) receptor tyrosine kinase ^14^, while its co-receptor glial cell-derived neurotrophic factor receptor alpha 1 (GFRA1) is expressed more broadly (**Fig. 1A,B** and **Fig. S1**). SIX2+ ‘cap mesenchymes’ of nephron progenitor cells and FOXD1+ stromal progenitors separating these caps signal to tip cells through secreted glial cell-derived neurotrophic factor (GDNF) ligand ^15–18^. Lineage-tracing experiments reveal a striking homing behavior of RET+ cells, where they cluster at nascent tips and remain throughout repeated branching cycles ^19^. GDNF-RET signaling activates downstream pathways such as RAS-RAF-MEK-ERK ^20^ and PI3K-AKT ^21,22^ that are required for normal branching ^23^.

**Figure 1.**
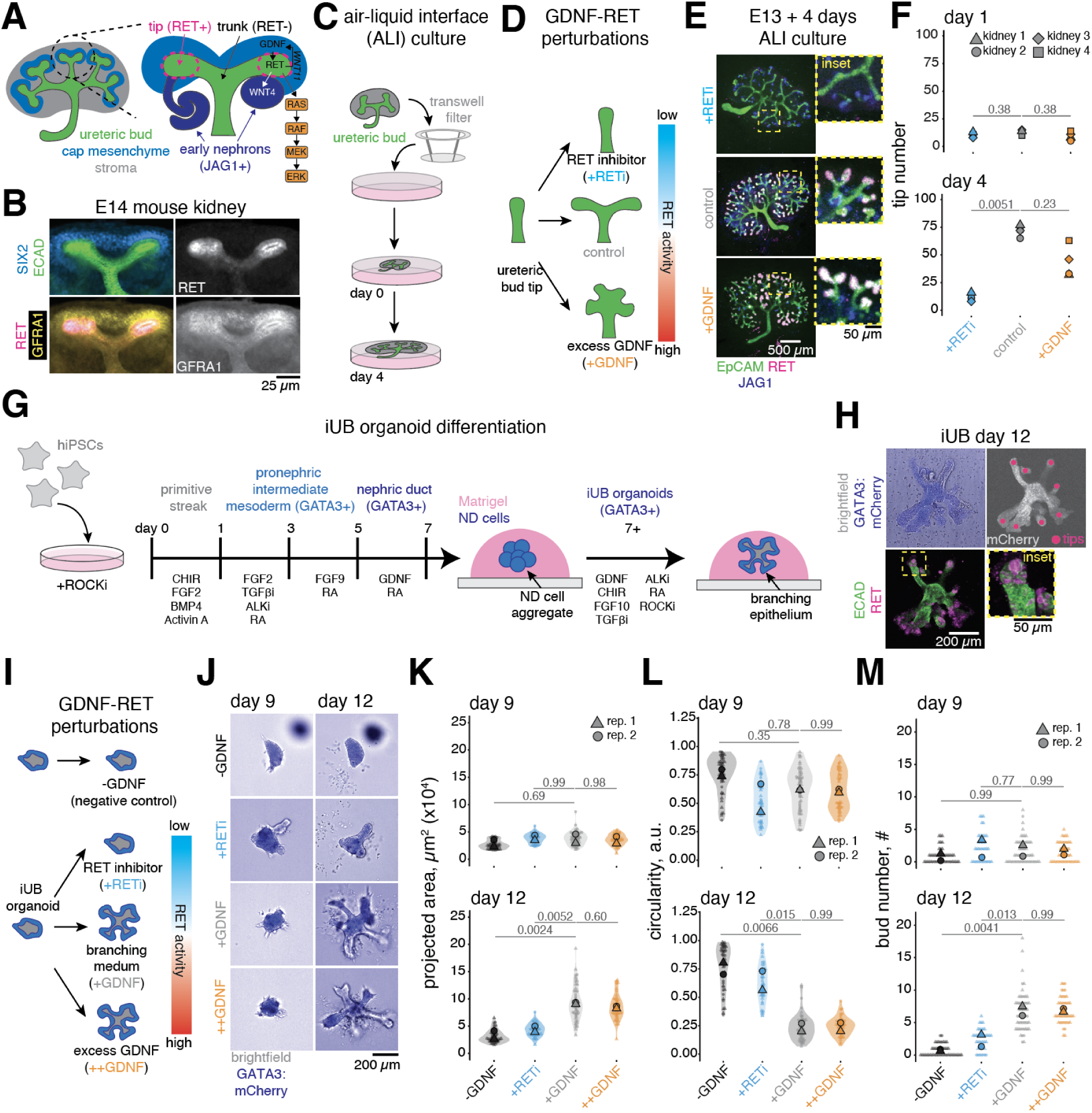
GDNF-RET signaling tunes branching in mouse and human ureteric bud tissues. A. GDNF-RET signaling interactions involved in kidney branching. B. Immunofluorescence of cleared E14 mouse kidney. *Top left*, ECAD and SIX2. *Bottom left,* RET and GFRA1. *Top right*, isolated RET. *Bottom right*, isolated GFRA1 signal. C. Air-liquid interface (ALI) culture of mouse embryonic kidney explants. D. Design of GDNF-RET signaling perturbation experiments in ALI culture. E. Immunofluorescence of E13 kidneys grown in ALI culture for 4 days in 100 nM Selpercatinib (+RETi), 100 ng ml^-1^ GDNF (+GDNF), or control media. *Inset*, close-up of tip domains. Explants are immunostained for EpCAM (epithelium), RET (tip cells), and JAG1 (early nephrons). F. Tip number on days 1-4 across all conditions, N = 4 kidneys per condition. *P*-values by one-way Kruskal-Wallis test with Dunn’s *post hoc* test. G. Differentiation of iUB organoids from hiPSCs H. Immunofluorescence of epithelial (ECAD) and tip (RET) markers in a day 12 iUB organoid. I. Design of GDNF-RET signaling perturbations for iUB organoids. J. Merged brightfield and GATA3:mCherry images of iUB organoids at days 9 and 12. Organoids were cultured in control (-GDNF) or complete branching medium (+GDNF, 50 ng ml^-1^), excess GDNF (++GDNF, 250 ng ml^-1^), or complete branching medium with 100 nM Selpercatinib (+RETi). See: **Fig. S4F**. K. Projected area (x10^4^ µm^2^) on days 9 and 12 for all conditions, n = 47, 56, 40, 46 iUB organoids (-GDNF, +RETi, +GDNF, ++GDNF) from 2 independent biological replicates. L. Circularity (a.u.) on days 9 and 12. M. Bud number on days 9 and 12. *P*-values in panels K, L, M by one-way ANOVA with Dunnett’s *post hoc* test using +GDNF as reference group.

Molecular genetics studies in mice have revealed that alterations to ERK signaling level within the ureteric bud can dramatically change branching outcomes ^24–26^, but ERK signaling alone is not sufficient to replace RET’s role in cell clustering and branching ^27^. This reflects a complex relationship between GDNF-RET signaling, tip cell identity, and cell movements during branching that is not fully understood.

Reconstructing the kidney’s *in vivo* manifold structure in organoids would be impactful for disease modeling studies and as tissue building blocks for artificial implantable kidneys ^28,29^. Human induced pluripotent stem cells (iPSCs) can be differentiated into induced ureteric bud epithelium (iUB) cell types that feature a tip-trunk hierarchy and rudimentary branching ^30–33^. These can then be recombined with nephron cell types ^34^ that integrate with iUB cells to achieve fluidic connectivity ^35^. However, state of the art iUB organoid protocols are presently limited to ∼1-3 branch generations with little control over branch location and frequency ^30^. Fully developed human kidneys have over 15 branch generations with stereotyped early branch events ^36,37^, posing a long-term design challenge. Since ureteric bud tips are the sites of nephron integration, limited branching extent currently hinders mosaic iUB:nephron organoids with controlled geometry and parallelization. Rather than relying on intrinsic self-organization of tip and trunk cells, optogenetic tools could specify signaling zones for bud initiation, elongation, and bifurcation with high spatial precision. This would permit construction of scalable ductal networks with defined architectures and nephron niches.

Here, we focus on achieving spatial control over bud initiation and elongation. We benchmark the relationship between GDNF-RET activity dosage, downstream signaling, and morphogenesis in mouse explants and human organoids. We then design and validate an optogenetic RET receptor (optoRET) to assert engineering control via blue light dosage. We find optoRET replicates quantitative control over ERK signaling and epithelial morphogenesis in mammalian cell lines and human iPSC-derived kidney organoids. Targeted optoRET stimulation by spatially patterned blue light guides morphogenesis in engineered iUB organoids in the absence of GDNF. Our results establish optoRET for interrogating developmental cell behaviors and for guided morphogenesis of synthetic tissues.

## Results

### GDNF-RET signaling levels determines branching outcomes in mouse and human kidney tissues

Ureteric bud branching in the embryo is acutely sensitive to genetic perturbation of GDNF-RET signaling ^38–41^, making this pathway an appealing control handle for organoid engineering.

However, the fact that RET, GDNF, or GFRA1 knockout leads to kidney agenesis with ∼90% penetrance in mice limits further study without complex conditional and tissue-specific genetic models ^38–41^. Mosaic models, where a fraction of ureteric bud cells are deficient in RET signaling ^14,42^ or express a constitutively active RET isoform ^27^, have revealed that RET activity is required for cell clustering, sorting into tips, and maintaining a tip:trunk hierarchy. To further explore how GDNF-RET signaling controls branching, we cultured E13 mouse kidneys in an air-liquid interface (ALI) culture system ^43^ that permits monitoring of tissue growth and branching morphogenesis *ex vivo* (**Fig 1C**). To perturb GDNF-RET signaling, we either added GDNF (+GDNF) or the RET-specific kinase inhibitor ^44^ Selpercatinib (+RETi) to the culture medium (**Fig. 1D**). ALI explants grew in area at similar rates across conditions over a 4 day period (**Fig. S2A,B**). Endpoint immunostaining revealed +GDNF kidneys had fewer tips than controls (**Fig. 1E,F**) but larger tip domains (**Fig. S2D,F**). By contrast, +RETi explants had fewer tips, did not form new terminal branches ^45^, and lacked detectable RET+ tip cell clusters (**Fig. 1E**). Early nephron structures (JAG1+) were present in all treatment conditions. Prior groups have noted a non-monotonic relationship between GDNF-RET signaling and tip number based on mouse genetic studies, where hypoactive or hyperactive RET signaling each lead to distinct kidney defects ^24,46,47^. Our *ex vivo* data suggest a ‘Goldilocks effect’ where an intermediate level of GDNF-RET signaling is required to support branching, while high or low deviations lead to distinct branching defects.

Next, we tested whether similar GDNF-RET perturbations would also tune bud formation and elongation in human iUB organoids ^30^. Here, iPSCs are differentiated into GATA3+ nephric duct (ND) spheroids over 7 days (**Fig. 1G, Fig. S3**, and **Methods**). ND spheroids are then transferred into 3D Matrigel droplets and cultured over days 7-12 in a ureteric bud medium (UBM) containing GDNF and other factors that support branching (**Table S1**). During this time, spheroids form paddle-like elongated buds and bifurcated branches that contain RET+ tip cells (**Fig. 1H**, **Fig. S3H,I**, and ref. ^30^). Live imaging revealed buds are dynamic, undergoing periods of net extension and retraction during organoid growth (**Fig. S4A-C** and **Movies S1, S2**). To further investigate morphological outcomes downstream of GDNF-RET, we compared organoids exposed to the standard GDNF concentration in UBM (+GDNF, 50 ng ml^-1^), UBM containing 5x higher GDNF (++GDNF, 250 ng ml^-1^), or UBM plus Selpercatinib (+RETi, 100 nM) starting at day 9 (**Fig. 1I-M**, **Fig. S4F**, **Table S1**, and **Methods**). Organoids in the ++GDNF group neither decreased in circularity nor increased in area or bud number compared to +GDNF organoids.

The latter observation contrasts with the ‘Goldilocks’ effect of GDNF-RET described for mouse kidney explants. Organoids in the +RETi condition remained small, circular in cross-section, and did not form new buds. As a negative control, we omitted GDNF over days 7-12 (-GDNF). Here we measured smaller projected areas, higher circularity, and fewer buds, as well as a reduction in the number of tip cells. Interestingly, we also found a linear scaling relationship between organoid area and bud number across the conditions (**Fig. S4G,H**). These results indicate similar morphogenetic plasticity in response to changes in GDNF-RET signaling between mouse and human ureteric bud tissues.

### GDNF-RET maintains a tip cell progenitor state whose correlation with ERK signaling depends on developmental context and signaling level

We next sought to comparatively assess the relationship between GDNF-RET and downstream ERK activity in mouse kidneys and human iUB organoids. A genetically encoded ERK biosensor previously revealed higher signaling activity in mouse tip vs. trunk cells ^20^, suggesting higher GDNF-RET activity there. MEK activity is also required for tip bifurcation in murine kidneys ^23,48^. We first immunostained E13 mouse kidneys grown in ALI culture in the same pharmacological conditions for RET and ppERK (**Fig. 2A** and **Fig. S5A,B**). Tip cells from +RETi kidneys had lower RET staining compared to controls, reflecting either a lower frequency of RET+ cell homing to tips ^27^ and/or lower positive feedback on RET expression ^49^ (**Fig. 2B,C**). ppERK signal was also lower, validating that GDNF-RET sustains ERK signaling in tip cells.

**Figure 2.**
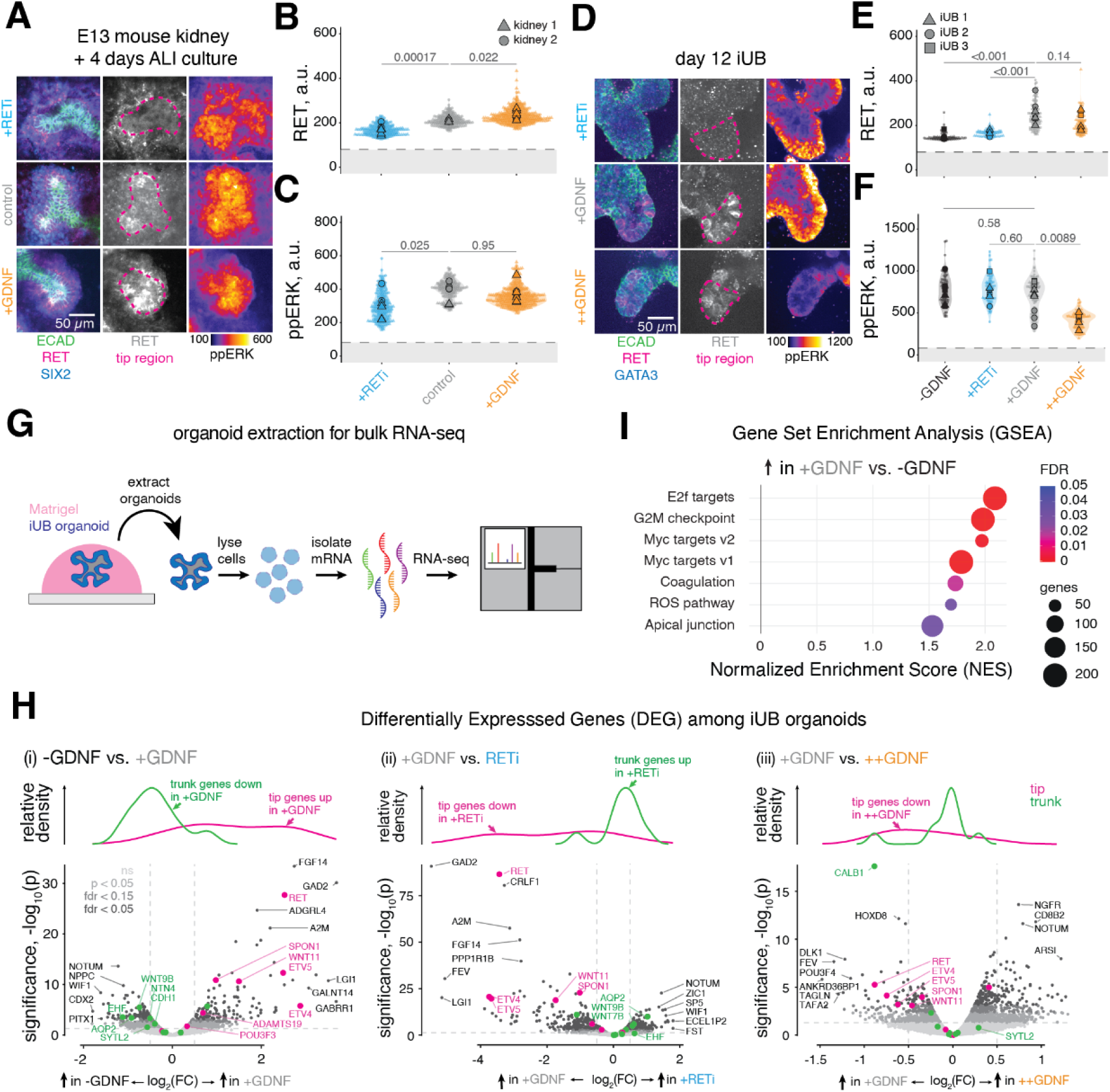
GDNF-RET sustains ERK signaling and tip identity among a tip progenitor population. A. Immunofluorescence of E13 kidneys grown in ALI culture for 4 days and treated with +RETi, +GDNF, or control, as described above. *Left*, epithelial (ECAD), ureteric bud tip (RET), and cap mesenchyme (SIX2) markers. *Middle*, isolated RET and tip outline. *Right*, intensity-coded ppErk (a.u.). B. Mean RET intensity (a.u.) of n = 197, 169, 312 cells measured from 6, 7, 8 tips (+RETi, control, +GDNF). C. Mean ppErk intensity (a.u.) for all tip cells. Data are pooled from 2 kidneys per condition. Dashed line in panels b, c shows microscope background cutoff (a.u.). D. Immunofluorescence of day 12 iUB organoids grown in -GDNF, +GDNF, ++GDNF, and +RETi conditions (see also: **Fig. S4F** and **Table S1**). *Left*, epithelial (ECAD) and tip cell (RET) markers. *Middle*, isolated RET and tip outline. *Right*, intensity-coded ppErk (a.u.). E. Mean RET intensity (a.u.) of n = 67, 72, 97, 81 cells measured from 7, 9, 7, 6 tips (-GDNF, +RETi, +GDNF, ++GDNF). F. Mean ppErk intensity (a.u.) for all tip cells. Data are pooled from 3 organoids per condition. Dashed line in panels e, f show the microscope background cutoff (a.u.). *P*-values in panels B, C, E, F by one-way ANOVA with Dunnett’s *post hoc* test using +GDNF as the reference group. G. Organoid preparation workflow for bulk RNA-seq. H. Differential expression analysis from bulk RNA-seq of day 12 iUB organoids grown in -GDNF, +GDNF, ++GDNF, and +RETi conditions (see also: **Fig. S6**). *Top row*, relative density of tip and trunk genes. *Bottom row*, volcano plots of all genes organized by significance (-log_10_(*p*)) and log_2_(fold-change) with tip and trunk-specific markers highlighted. *Left*, -GDNF vs. +GDNF groups. *Middle*, +GDNF vs. ++GDNF groups. *Right*, +GDNF vs. +RETi groups. Data are pooled from 3 biological replicates. I. Gene Set Enrichment Analysis (GSEA) of upregulated genes in day 12 iUB organoids between the +GDNF and -GDNF groups as a function of normalized enrichment score (NES). Hallmark gene sets are color coded by false discovery rate (FDR), size coded by the size of the gene set.

Unexpectedly, while +GDNF kidneys clearly had larger tips, the average ppERK intensity remained similar to controls (**Fig. 2B,C**). Our results are consistent with a model where maintenance of the tip cell compartment by homing or transcriptional positive feedback requires RET activity. However, while ERK signaling may be necessary ^50^ it is not sufficient for this, at least at super-physiological activity levels. This matches earlier conclusions from a mouse model with mosaic expression of constitutively active MEK ^27^.

We next used the same pharmacological manipulations of GDNF-RET to examine RET and ppERK in day 12 iUB organoids for which far less is known. We found RET+ cells were present within the distal ∼50-100 µm of branch tips in the +GDNF condition (**Fig. S3H,I**) so we compared the RET and ppERK intensities among these most distal bud cells across all conditions (**Fig. 2D, Fig. S5C,D**). RET intensities (**Fig. 2E**) followed an expected trend we had observed in mouse kidney explants (**Fig. 2B**), where the -GDNF and +RETi samples had lower RET signal than +GDNF or ++GDNF samples. This suggests a reduction in the number of tip progenitor cells under reduced GDNF-RET signaling conditions. Intriguingly, ppERK was similar between -GDNF, RETi, and +GDNF treatment groups (**Fig. 2F**), suggesting that RET is not the sole activator of ERK signaling in iUB organoid tip cells. Such a discrepancy could be explained by the presence of other RTK ligands (such as FGF10, ref. ^25^) in the culture medium (**Table S1**), or by physical mechanisms such as curvature-mediated regulation of ERK (ref. ^8^). These data show that iUB organoids have an intact positive feedback loop on tip cell state mediated by GDNF-RET, but where the downstream role of ERK may be obscured by redundancy specific to organoid culture context.

To further characterize this positive feedback, we performed bulk RNA-seq and Differential Gene Expression (DEG) analysis (**Fig. 2G-H, Fig. S6**) and Gene Set Enrichment Analysis (GSEA) (**Fig. 2I**). For DEG, we curated tip and trunk gene sets by combining known markers from our previous study on week 20 human embryonic kidney spatial sequencing data with hits from a screen for tip vs. trunk markers in E15.5 mouse kidneys ^13,17^. Expectedly ^49^, the +GDNF condition showed enrichment for tip genes and depletion of trunk gene expression relative to -GDNF and RETi conditions. The ++GDNF condition reduced rather than increased tip gene expression relative to +GDNF, curiously similar to the ‘Goldilocks’ effect we observed in mouse explants (**Fig. 1F**). Gene sets enriched in +GDNF vs. -GDNF conditions were dominated by those associated with cell proliferation (G2M checkpoint, E2F targets, MYC targets) and tracked with tip vs. trunk gene expression changes (**Fig. 2I**). These data indicate that changes in GDNF-RET signaling manifest in *bona fide* manipulation of tip vs. trunk cell composition and not solely spatial redistribution of existing RET+ cells by homing to tips. Moreover, the data add to our other evidence that the GDNF-RET axis has a similar effect in maintaining tip cell identity in human iUB organoids as in the embryonic mouse kidney.

### An optoRET receptor controls ERK-dependent epithelial morphogenesis

Our explant and organoid results validated the GDNF-RET axis as a viable target for synthetic morphogenesis studies using optoRTKs. Established optoRTK designs replace the ligand-selective extracellular domain with an intracellular light-sensitive protein domain ^11^, while physiologic RET signaling requires the formation of a tripartite complex with GDNF and GFRA1^15^. As a benchmark for optogenetic tool sensitivity, we generated HEK 293T cells co-expressing RET and GFRA1 and measured a robust increase in ppERK immunofluorescence after stimulation with GDNF (**Fig. S7**). We then designed an optogenetic RET (optoRET) receptor (**Methods**) containing the human RET9 intracellular domain (ICD) flanked by an N-terminal myristoylation peptide for membrane insertion, and a C-terminal module containing a red fluorescent protein (mCherry) and the *A. thaliana* Cryptochrome 2 photolyase homology region (CRY2^PHR^) as a blue light-sensitive actuator (refs. ^12,51^) (**Fig. 3A**). HEK 293T cell lines expressing optoRET (HEK-optoRET cells) were produced by viral induction (**Fig. 3B**), purified by fluorescence activated cell sorting (FACS), and stimulated with 470 nm light (0-160 mW cm^-2^, 1 s every 10 s) for 2 hrs using the LED array on an optoPlate-96 ^52^ to evaluate their sensitivity (**Fig. 3C**). Quantitative immunofluorescence analysis of ppERK revealed a similar dynamic range to ligand stimulation (∼10-fold increase), with saturation above ∼16 mW cm^-2^ and half-maximal activation ∼6 mW cm^-2^ (**Fig. S8** and **Methods**). This indicates blue light stimulation of optoRET induces downstream ERK signaling over an appropriate physiological range.

**Figure 3.**
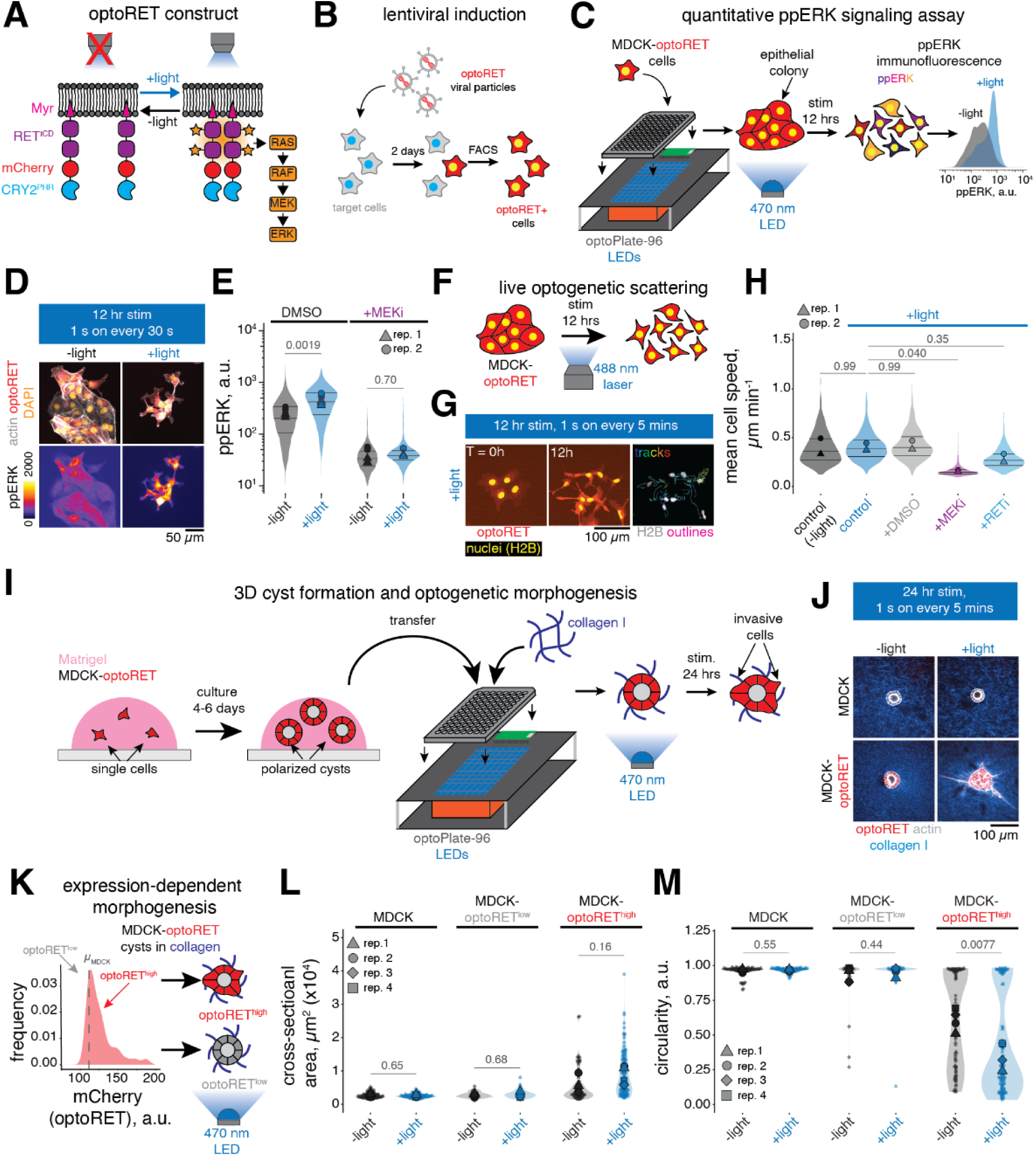
Blue light stimulation of optoRET drives ERK-dependent symmetry breaking in epithelial cysts. A. OptoRET design consisting of a Myristoylation domain (Myr), RET intracellular domain (RET^ICD^), red fluorescent protein (mCherry), and *A. thaliana* Cryptochrome 2 photolyase homology region (CRY2^PHR^). Stimulation with 470 nm light causes CRY2^PHR^ oligomerization and signaling through RAS-RAF-MEK-ERK. B. Viral production and generation of stable optoRET-expressing cell lines. C. Schematic for blue light stimulation of MDCK-optoRET cells in the optoPlate-96 and density plot of ppERK (a.u.) for n = 987, 1082 cells after 12 hr stimulation with 470 nm light (+light, 50 mW cm^-2^, 1 s every 30 s) or left unstimulated (-light). D. Immunofluorescence of MDCK-optoRET cells following 12 hrs stimulation with 470 nm light (50 mW cm^-2^, 1 s every 30 s). *Top*, actin (phalloidin), optoRET (mCherry), and DAPI. *Bottom*, intensity-coded ppERK (a.u.) for all conditions. E. Mean ppERK (a.u.) measured in MDCK-optoRET cells for the above stimulation conditions and treated with either DMSO or 100 nM Trametinib (+MEKi) for the duration of stimulation. DMSO: n = 3181, 2926 cells, +MEKi: n = 3210, 2086 cells (-light, +light), pooled from 2 biological replicates. *P*-values by Welch’s t-test. F. Live MDCK-optoRET scattering by 488 nm confocal stimulation. G. Example image sequence of MDCK-optoRET cells co-expressing H2B-mVenus at T = 0 and 12 hrs and example TrackMate ^56^ output overlay. Cells were imaged once every 5 mins and stimulated with the 488 nm laser (220 mW cm^-2^, 1 s every 5 min). H. Mean cell speeds obtained by single cell tracking of MDCK-optoRET cells. Cells were untreated (-light, +light) or treated with DMSO, 100 nM Trametinib (+MEKi), or 100 nM Selpercatinib (+RETi) for the duration of imaging, n = 348, 414, 520, 677, 574, cells (-light, +light, +DMSO, +MEKi, +RETi) pooled from 2 biological replicates. All conditions except negative controls (-light) were stimulated as described above. *P*-values by one-way ANOVA with Dunnett’s *post hoc* test. I. Procedure for MDCK cyst formation in Matrigel, transfer to collagen I gels, and morphogenesis assay using the optoPlate-96. Cysts were transferred to collagen I gels and were stimulated for 24 hrs with 470 nm light. J. Immunofluorescence of MDCK and MDCK-optoRET cysts in a 2 mg ml^-1^ collagen I gel following 24 hrs stimulation with 470 nm light at 0 mW cm^-2^ (-light) or 50 mW cm^-2^ (+light). Collagen I fibers were pre-labeled with Alexa 647-NHS and actin was visualized using phalloidin. K. Data filtering scheme for MDCK-optoRET cysts. Samples were binned into optoRET^high^ and optoRET^low^ groups by average mCherry intensity (a.u.) of MDCK cysts (µ_MDCK_). OptoRET^high^ and optoRET^low^ cyst groups were binned and analyzed separately (see also: **Fig. S11A** and **Methods**). L. Cross-sectional area (µm^2^ x10^4^) of MDCK and MDCK-optoRET cysts, n = 93, 95 MDCK cysts, n = 32, 36 optoRET^low^ cysts, and n = 83, 115 optoRET^high^ cysts (-light, +light) pooled from 4 biological replicates. M. Circularity (a.u.) of cysts in panel k. *P*-values in panels K, L by Welch’s t-test.

Madin-Darby Canine Kidney (MDCK) epithelial cells are a better model system for connecting RTK signaling to cell collective behaviors ^53^. MDCKs expressing full-length human RET (MDCK-RET cells ^54^), but not a kinase-dead RET mutant (MDCK-KM cells), activate ERK ^22^ in response to stimulation with GDNF and soluble GFRA1. This in turn drives cell scattering (**Fig. S9A-G**, **Movie S3**). To test the capacity of optoRET to reproduce these behaviors, we produced stable MDCK-optoRET cell lines by lentiviral transduction and FACS (**Methods**). To evaluate ERK signaling responses, we stimulated cells with 470 nm light (50 mW cm^-2^, 1 s every 30 s) for 12 hr using the optoPlate-96 (**Fig. 3C,D**). Light-stimulated cells formed long, actin-based protrusions, dispersed from colonies, and showed higher ppERK than unstimulated controls (**Fig. 3D,E**). Concurrent treatment with 100 nM Trametinib (MEKi) abrogated the ppERK increase and caused cells to remain in tight colonies (**Fig. 3E**). We confirmed live scattering responses of MDCK-optoRET to light time lapse imaging over 12 hrs with concurrent stimulation provided by a 488 nm laser line (**Fig. 3F, G**). Surprisingly, semi-automated nuclear tracking revealed that the protrusion and scattering dynamics downstream of optoRET did not involve an increase in cell speed compared to unstimulated controls (**Fig. 3H**, **Fig. S10**, **Movie S4**), suggesting other mechanisms such as downregulation of cell-cell adhesion ^54^. Concurrent treatment with 100 nM Trametinib (+MEKi), or 100 nM Selpercatinib (+RETi) dramatically reduced cell speeds and abolished scattering. These data indicate optoRET stimulation replicates a RET- and ERK-dependent MDCK cell scattering behavior.

MDCK cysts embedded in collagen gels break symmetry and form invasive fronts in a well-documented response to RTK ligands ^53–55^. We adapted a previously described cyst morphogenesis assay ^55^ to test whether blue light stimulation of optoRET could cause similar symmetry breaking. Single cells embedded in Matrigel formed cysts over 4-6 days that were then transferred to collagen I gels (**Fig. 3I**). We first prototyped the assay using the MDCK-RET cells. Following GDNF stimulation, cells with higher RET expression invaded the surrounding gel as single cells or multicellular chains with a leader-follower organization (**Fig. S9H,I**, **Movie S5**). This led to an increase in midplane area and loss of circularity in a fraction of cysts within ∼48 hr of stimulation with GDNF (**Fig. S9J,K**).

Moving to the MDCK-optoRET cells, we exposed cysts to 470 nm light (50 mW cm^-2^, 1 s every 5 mins) using the optoPlate-96 (**Fig. 3I**). Light-stimulated cysts broke symmetry and began to invade the surrounding collagen matrix (**Fig. 3J**, **Movie S6**). We also observed a sub-population of un-stimulated cysts that were larger and exhibited long protrusions (decreased circularity), consistent with autoactivation in the absence of light. Since the most invasive cysts appeared to have higher optoRET (mCherry) expression, we filtered cells into an optoRET^high^ and optoRET^low^ population based on the mean plus standard deviation mCherry signal of optoRET- MDCK cysts (µ_MDCK_, **Fig. 3K**, **Fig. S11A,** and **Methods**). Stimulated optoRET^high^ cysts (+light) had larger midplane areas (**Fig. 3L** and **Fig. S11B**) and smaller circularities (**Fig. 3M** and **Fig. S11C**) within ∼24 hr of stimulation than unstimulated cysts (-light). Cysts with sub-threshold expression (optoRET^low^) showed neither morphogenesis among unstimulated controls, nor responsiveness to light. In a separate experiment, concurrent treatment of the optoRET^high^ group with MEKi completely blocked light-induced morphogenesis compared to vehicle (DMSO) controls (**Fig. S11D-F**). The data suggest optoRET expression level is an important parameter in controlled morphogenesis, where expression must be high enough such that light-induced optoRET activation is sufficient to trigger morphological change, but not so high that autoactivation dominates.

### Spatially patterned optoRET stimulation controls human kidney organoid budding

We next sought to test whether we could instruct branching outcomes with blue light in a human epithelial tissue. We generated human iPSC lines expressing optoRET using PiggyBac (hiPSC-optoRET, **Fig. 4A**, **Methods**, **Movie S7**) and differentiated them into ND spheroids (ND-optoRET spheroids) using the 7-day differentiation protocol (**Fig. 4A, Methods**). OptoRET integration did not disrupt normal iUB lineage commitment in organoids (iUB-optoRET organoids), as confirmed by a GATA3 endogenous reporter ^57^ and PAX2 and RET immunofluorescence (**Fig. S12**).

**Figure 4.**
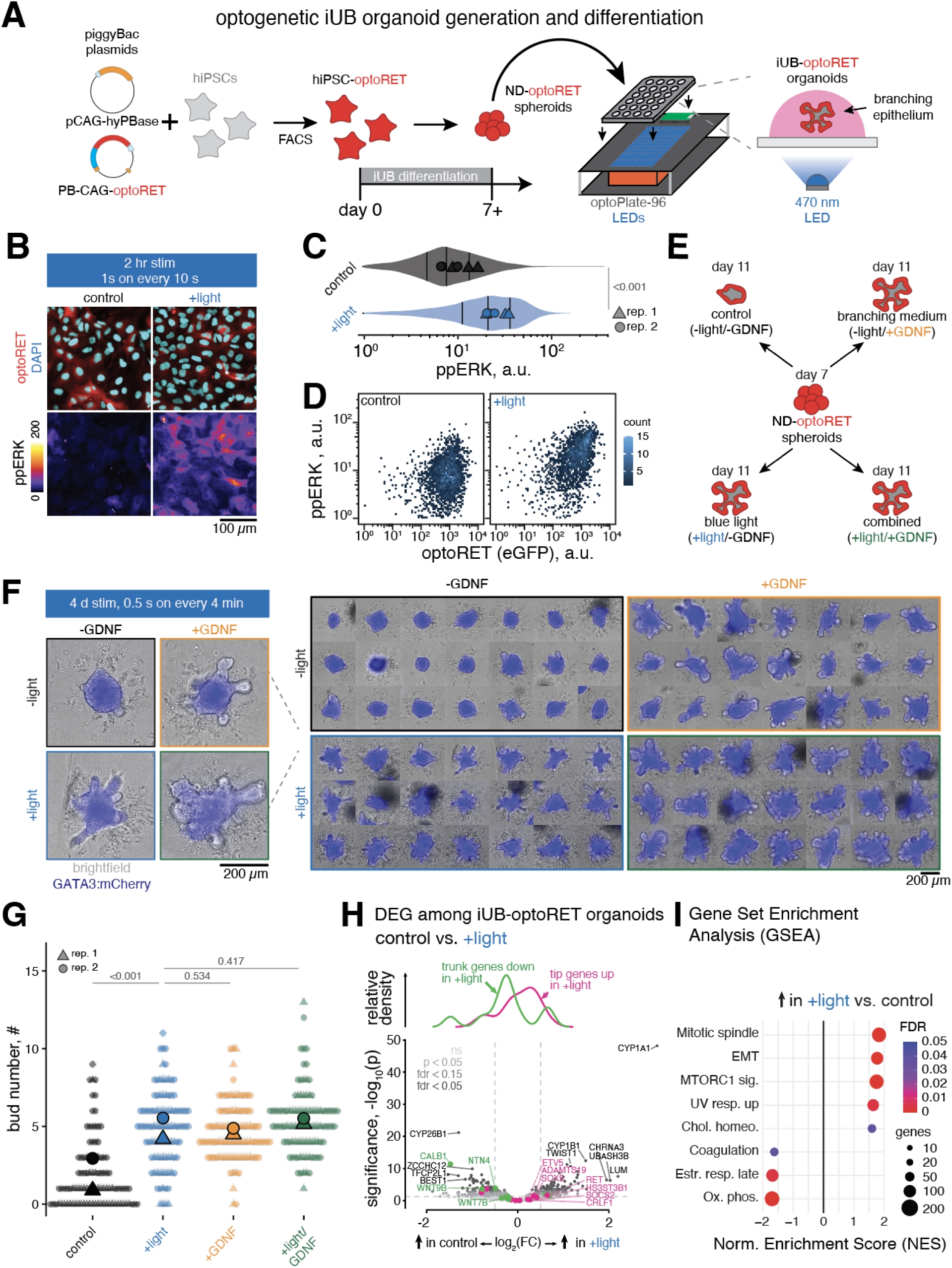
Blue light stimulation of optoRET drives ERK signaling and ligand-independent budding in iUB organoids. C. Schematic of hiPSC-optoRET generation using piggyBac transposase and differentiation into iUB-optoRET organoids. D. Immunofluorescence of iUB-optoRET monolayers stimulated for 2 hr under the indicated conditions (±light). Blue light stimulation was provided by an optoPlate-96 (470 nm, 50 mW cm^-2^, 1 s every 10 s) for 2 hrs. *Top*, optoRET (detected by anti-GFP) and nuclei (DAPI). *Bottom*, intensity-coded ppERK (a.u.). E. Violin plot of ppERK (a.u.) for iUB-optoRET cells under -light and +light conditions, n = 2458, 2030 cells (control, +light) pooled from 2 independent biological replicates. *P*-value by Welch’s t-test. F. Immunofluorescence density plot of ppERK (a.u.) as a function of optoRET(eGFP) intensity (a.u.) for cells shown in B. n = 2458, 2030 cells (control, +light) pooled from two independent biological replicates. G. Stimulation conditions for iUB-optoRET organoids: control (−light/−GDNF), +light, +GDNF, and combined (+light/+GDNF) treatments. H. Live fluorescence images of GATA3^mCherry^ iUB-optoRET organoids under optogenetic and ligand-based stimulation conditions. Blue light stimulation was provided by an optoPlate-96 (320 mW cm^-2^, 0.5 s every 4 min) between days 7-11 and the +GDNF and +light/+GDNF groups received 50 ng ml^-1^ GDNF. I. Bud number on day 11 across all conditions, n = 50, 50, 51, 50 organoids (control, +light, +GDNF, +light/+GDNF) from 2 independent biological replicates. *P*-values by one-way Kruskal-Wallis test with Dunn’s *post hoc* test. J. Differential expression analysis from bulk RNA-seq of iUB-optoRET organoids grown in ±light stimulation conditions (see also: **Fig. S14**). *Top*, relative density of tip and trunk marker genes. *Bottom*, volcano plot of all genes organized by significance (-log_10_(*p*)) and log_2_(fold-change) with tip and trunk-specific markers highlighted. Data are derived from 3 biological replicates. K. Gene Set Enrichment Analysis (GSEA) as a function of normalized enrichment score (NES). Hallmark gene sets are color coded by false discovery rate (FDR), size coded by the size of the gene set.

To determine whether optoRET activation could stimulate ERK signaling among iUB cells, we seeded day 7 ND-optoRET spheroids into Matrigel-coated 96-well plates and allowed them to spread into monolayers overnight (**Methods**). The monolayers were then stimulated for 2 hr on an optoPlate-96 under control (-light/-GDNF) and blue light (+light/-GDNF) conditions (**Methods**). The blue light condition increased ppERK across iUB-optoRET monolayers (**Fig. 4B,C**), while the control condition maintained low baseline ppERK levels as expected. Consistent with our data in HEK cells (**Fig. S8A**), higher optoRET level in a given cell correlated with increased ppERK intensity in that cell following light stimulation (**Fig. 4D**). These data confirm that iUB-optoRET cells are functionally responsive to optogenetic stimulation.

We next asked whether blue light stimulation could direct ligand-free branch initiation or elongation in iUB-optoRET organoids. We plated day 7 ND-optoRET spheroids in Matrigel and cultured them under control (-light/-GDNF), blue light (+light/-GDNF), branching medium (-light/+GDNF), or combined (+light/+GDNF) conditions (**Fig. 4E, Methods**). iUB-optoRET organoids in the blue light or combined conditions were stimulated with uniform 470 nm blue light (320 mW cm^-2^, 0.5 s every 4 min) from days 7-11 using the optoPlate-96 (**Fig. 4F**). Blue light alone significantly increased budding events relative to unstimulated controls (**Fig. 4G**), demonstrating that optoRET activation is sufficient to initiate budding in the absence of exogenous GDNF. Similar results were obtained in an independent hiPSC cell line (**Fig. S13**). Bulk RNA sequencing of light-stimulated organoids indicated a mild increase in tip marker genes at the expense of trunk markers and an increase in cell proliferation-related gene sets relative to unstimulated controls (**Fig. 4H,I; Fig. S14**). Though muted, both effects were similar to the corresponding +GDNF vs. control comparisons (**Fig. S14**), indicating that stimulating optoRET has qualitatively similar effects on tip vs. trunk cell composition as the endogenous GDNF-RET axis (**Fig. 2H,I** and ref. ^49^). A lack of quantitative correspondence may relate to a lower engagement of optoRET with positive transcriptional feedback on tip cell state, or a variety of technical factors including the light stimulation program, optoRET transduction efficiency, etc. A generic epithelial-to-mesenchymal transition (EMT) gene set was also enriched in +light but not +GDNF conditions. Together the data show successful reconstitution of the morphological and molecular effects of GDNF-RET signaling by optoRET in iUB organoids.

Geometric control of stimulation is a powerful advantage of optogenetics over diffusible ligand-based stimulation, since light can be readily delivered to tissues with subcellular precision. Therefore, we asked whether spatially patterned optoRET stimulation could instruct the locations of new branch initiation and produce asymmetric branching outcomes in iUB organoids. To test this, we used a spinning disk microscope equipped with a digital micromirror device (DMD) and 488 nm LED illumination (**Methods**). DMD intensity was significantly higher than that from the opto-Plate-96 in a side-by-side fluorescent bead-based photobleaching calibration (**Fig. S15**). We aligned a stimulation region of interest (ROI) either across the entire organoid (+light whole) or asymmetrically, across the right-hand half of the organoid (+light half) within the imaging field (**Fig. 5A**). Unstimulated organoids served as negative controls and we included a +GDNF group as a positive control, since we previously noted these had no apparent branching asymmetry (**Fig. 1J**). We seeded organoids at two different initial diameters (⌀_small_ = 174 ± 13 µm, and ⌀_large_ = 284 ± 41 µm, mean ± S.D., **Fig. 5B,C**) and found that stimulating entire organoids (+light whole) caused the formation of new buds throughout the illuminated ROI over 4 days (**Movie S8**). Under DMD stimulation, new tubules induced in the light-exposed (+light whole and +light half) groups were smaller in diameter than ones in the +GDNF group. A ‘skirt’ of invading cells was more prominent throughout the stimulated area, most of which expressed optoRET at high levels and lacked ECAD expression (**Fig. 5D, S12E**). These solitary invading cells are in line with the generic EMT signature in optoRET-stimulated organoid bulk RNA-seq results (**Fig. 4I**). Immunostaining for endogenous RET+ cells using an antibody raised against the RET extracellular domain (ECD) revealed tips containing intermingled cells expressing RET, optoRET, or both (**Fig. 5D**). This indicates normal RET+ cell recruitment and/or renewal at tips, whether those cells also expressed optoRET or not. A transition to full cell-autonomous invasion by individual optoRET+ trunk cells may have been suppressed by the fact that activated cells can also sort to bud tips, whether recruited through endogenous GDNF-RET or optoRET (**Fig. S3H,I** and **Fig. S12E**). This would create a counteracting ‘winner take all’ effect that consolidates the number of invasion sites and therefore bud number ^27^.

**Figure 5.**
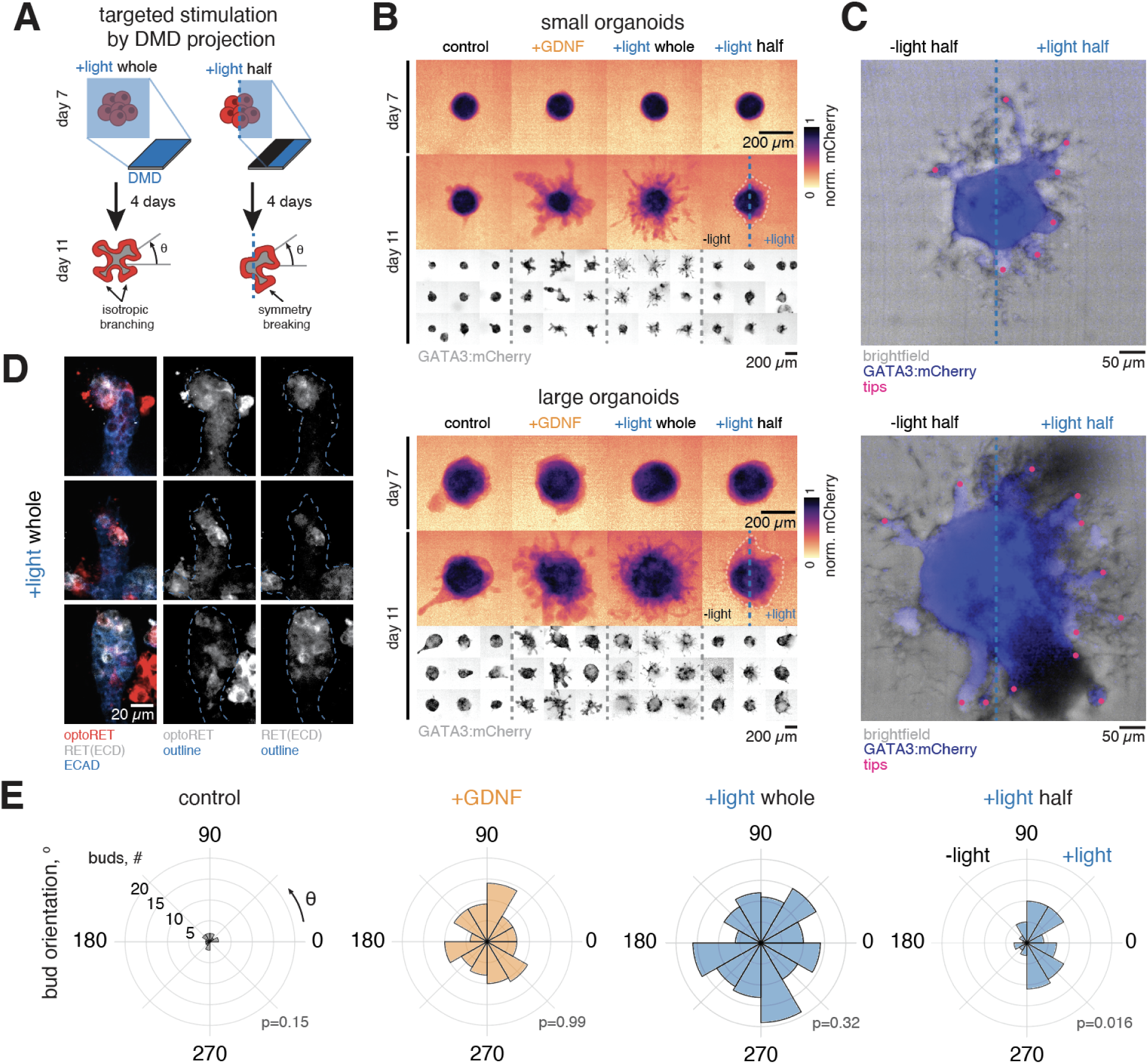
Spatially patterned optoRET stimulation drives asymmetric budding in iUB organoids. A. Strategy for targeted optogenetic stimulation of iUB-optoRET organoids. B. Average projections of GATA3:mCherry for small (*top panels*) and large (*bottom panels*) iUB-optoRET organoids under control, +GDNF, +light whole, and +light half conditions. Light stimulation was provided by 488 nm DMD-based light projection (0.5 s every 4 mins) for 4 days. Organoids in the +GDNF group received 50 ng ml^-1^ GDNF. *Top row*, average projections across individual organoids at day 7. *Middle row*, average projections across individual organoids at day 11. *Bottom row*, normalized average projections for individual organoids at day 11. C. Representative average projections of individual small (*top*) and large (*bottom*) iUB-optoRET organoids on day 11. The cyan dashed line divides the non-illuminated side (*left*, -light half) from the illuminated side (*right*, +light half). Magenta dots indicate bud locations. D. Confocal immunofluorescence images of optoRET+ and RET+ cells within iUB-optoRET bud tips in +light whole condition. Endogenous RET was visualized with an antibody against the extracellular domain (ECD). *Left*, ECAD, optoRET, and RET(ECD). *Middle*, optoRET and tip outline. *Right*, RET(ECD) and tip outline. E. Radial histograms of bud orientation angle (°) for small organoids across all conditions, n = 19, 20, 19, 20 organoids (control, +GDNF, +light whole, and +light half) pooled from 2 biological replicates. *P*-values by Kolmogorov-Smirnov test against a uniform reference distribution.

Competition between single cell invasion, new bud initiation, and recruitment to existing buds may be adjusted by tuning light exposure parameters, optoRET expression level, and the ratio of optoRET+ to optoRET- cells in mosaic organoids in future work. The data together support a model where optoRET+ cells can form new buds with tip domains that have an organotypic enrichment of RET+ cells.

Under asymmetric (+light half) stimulation, organoids formed new buds with a directional bias toward the illuminated side (p = 0.016, Kolmogorov-Smirnov test against a uniform reference distribution, **Fig. 5E, Movie S8,** and **Methods**). By contrast, none of the other groups were statistically different from a uniform distribution (control: *p* = 0.15, +GDNF: *p* = 0.99, +light whole: *p* = 0.32, Kolmogorov-Smirnov test) suggesting they did not produce a directional bias in bud orientation. Intriguingly, among buds in the asymmetric (+light half) stimulation group, there was also a bias in tubule outgrowth tangent to the illuminated zone, recalling the orthogonal orientation of tubule branching at the kidney surface relative to the corticomedullary (shallow-to-deep) gradient of GDNF ^16^. These results demonstrate that spatially patterned activation of optoRET is sufficient to control the locations of bud initiation in iUB organoids. They also constitute proof-of-principle data for ‘remote control’ over the orientation of tubule outgrowth in kidney organoids that may translate to other epithelial organoid systems.

## Discussion

Kidney branching morphogenesis is an amalgamation of multiple coordinated cellular behaviors. We find that optoRET successfully reproduces several of these, including positive feedback on tip cell state ^49,58^, cell homing to tips ^19^, clustering ^27^, and proliferation ^59^. Blue light activation of optoRET faithfully reproduces stereotyped signaling and epithelial morphogenesis behaviors in cell culture, as well as ligand-free budding and branching in kidney epithelial organoids. We further demonstrate that spatial patterning of light cues can instruct asymmetric iUB organoid budding outcomes that would otherwise be challenging to achieve with diffusible ligands.

Orthogonal control of organoid morphogenesis through optogenetics opens new opportunities for fundamental insights into RET biology in the developing kidney and new epithelial organoid biomanufacturing approaches.

Known mutations in GDNF, RET, and GFRA1 ^60,61^ account for ∼5% of cases of monogenic kidney anomalies in human patients, motivating future studies that leverage optoRET to model understudied congenital kidney diseases. Similarly, optoRET could be used to parse how RAS-RAF-MEK-ERK, PI3K-AKT, and PKC ^62^ mediate cell behaviors, tip cell identity, or morphological selectivity downstream of RET. For example, recent evidence suggests MEK signaling tunes tip cell adhesion and volume fluctuations to favor branching ^48^. Yet, we found that although MEK-ERK signaling was an appropriate indicator of RET activity across cell contexts, it did not correlate with RET activity and was therefore not specific to tip cell state. Future studies could combine optoRET with live signaling reporters to decipher the role of RET within a larger branching regulatory network. This may shed light on the contributions of factors not directly controlled by optoRET, such as GFRA1, Sprouty (Spry1), and FGF10 ^25^. In this way, optoRET could be a powerful tool in resolving a long-standing gap in understanding how cell signaling coordinates biophysical properties during tubule budding, elongation, and bifurcation.

OptoRET offers an exciting opportunity to study the cell non-autonomous interactions with nephron progenitors that interact with tips *in vivo*. This feedback is thought to be crucial for both ureteric bud tip and nephron progenitor renewal vs. differentiation decision-making ^63,64^.

OptoRET could be employed to understand if RET pathway signaling strength, periodicity, and effect on RET+ cell homing to the tip have commensurate or divergent effects on nephron progenitor recruitment, adhesion/motility, and differentiation state local to activated tips. These experiments could be performed in an iPSC-derived mosaic organoid setting where addition of GDNF would not give sufficient cell type specificity.

From a forward-engineering standpoint, optogenetics is well-suited to construct branching networks at physiological scale. Light can be projected onto organoids with micrometer spatial resolution using a variety of techniques (DMD, laser scanning, holography) and light stimuli can be added or removed nearly instantaneously to mediate signal strength. Similar photopatterning approaches have relied on photodegradable biomaterials to instruct the location of budding structures in intestinal organoids ^65^, while our results indicate bud location can also be directed by mimicking developmental RTK signaling. Our findings therefore inspire new opportunities for ‘bottom up’ optogenetic manufacturing of branched epithelial building blocks for mosaic organoid construction. An epithelial tissue with pre-patterned niches could specify sites for nephron progenitor cell integration ^35^. OptoRET-expressing cells could also be combined with other biofabrication techniques (e.g. bioprinting, micromolding) to increase the reproducibility of branched networks. For example, a bioprinted structure mimicking early ureteric bud bifurcations ^66^ could be used to template subsequent bud locations under optogenetic control.

Further opportunities are found in engineering other complex epithelia, such as the lung, salivary gland, or pancreas. While branching morphogenesis mechanisms appear to be organ specific ^1^, many feature RTK signaling among a specialized population of tip cells that are similar in transcriptional state ^13^. OptoRTKs could therefore be designed and implemented in iPSCs to control tissue-specific branching dynamics guided by *a priori* knowledge of endogenous RTK signaling dynamics. An RTK-agnostic approach could also reveal the contributions of specific downstream signals by mixing and matching between organ systems. Our results in mouse tissues, cell lines, and human organoids therefore establish a generic strategy for synthetic control of branching morphogenesis.

## Study limitations

Limitations of this study motivate future work to determine if optoRET recapitulates the full range of proposed RET-induced behaviors among ureteric bud tip cells, especially in converting trunk cells to tip cells, homing/sorting of activated cells to nascent tips, achieving positive feedback on tip cell state, and closing the reciprocal feedback loop with adjacent nephron progenitors. These factors are important determinants of whether appropriate optoRET activity will reconstitute *in vivo*-like persistent, self-sustaining nephrogenic niches in mosaic organoids. Several of these areas will require a more nuanced approach to distinguishing between native RET and engineered optoRET signaling within the same cell and among neighboring cells. This study also makes preliminary progress toward understanding selectivity of RET-active ureteric bud tips among budding, tubule elongation, and branching behaviors. Creating further understanding of the underlying signaling inputs and mapping them to biophysical properties, guided by physics-based modeling, would be illuminating here. More broadly, it remains to be shown whether similar single-axis optogenetic control strategies will be equally effective in other epithelial organogenesis contexts.

## Acknowledgements

We thank members of the Hughes, Bugaj, and McCracken laboratories for helpful comments, advice, and experimental assistance, particularly Catherine Porter and Emma Warrner for assistance with iPSC culture and differentiation, Jiageng Liu for assistance with FACS, and David Gonzalez-Martinez and Lee Roth for advice on cloning and optogenetics. We thank Gregory Dressler (University of Michigan) for the gift of MDCK-RET and MDCK-KM cells, Lois Mulligan (Queen’s University) for the human RET9, RET51, and GFRA1 plasmids, Harold Janovjak (Flinders University) for the opto-hRET plasmid, and Wenli Yang and the Human Pluripotent Stem Cell Core (Children’s Hospital of Philadelphia) for assistance with iPSC culture and differentiation. Cell sorting was performed in part on a BD FACSAria Fusion obtained through NIH grant S10 1S10OD026986 (L.J.B. and A.J.H.) and on other instruments operated and maintained by the Penn Cytomics and Cell Sorting Resource Laboratory. This work was supported by NIH F32 fellowship DK126385 and a Penn Center for Soft & Living Matter fellowship (L.S.P.), an NIH PERFORM-KUH Training Program TL1-DK143326 and U2C-DK136784 award (D.S.A), the NSF Graduate Research Fellowship Program (W.B.), NSF CAREER award 2047271 (A.J.H.), NIH NIGMS MIRA awards R35GM133380 (A.J.H.) and R35GM138211 (L.J.B.), NIH NIDDK R01DK132296 (A.J.H.), a Penn Center for Precision Engineering for Health (CPE4H) pilot grant (A.J.H.), a Commonwealth Universal Research Enrichment (CURE) grant 585499 from the Pennsylvania Department of Health (A.J.H.), a Child Health Research Career Development Award NIH NICHD K12HD028827 (K.W.M.), and a Carl W. Gottschalk Research Scholar Grant from the American Society of Nephrology (K.W.M.). No federal funds were used to collect human kidney tissue for Xenium spatial sequencing.

## Author contributions

Conceptualization, L.S.P. and A.J.H.

Methodology, L.S.P., R.C., A.Z.H., D.S.A., S.L.S., W.B., S.H.G., Z.D.H., T.R.M., and A.J.H. Software, L.S.P., R.C., S.L.S., W.B., L.J.B., and A.J.H.

Validation, L.S.P., R.C., A.Z.H., S.L.S., and D.S.A.

Formal analysis, L.S.P., R.C., and A.J.H. Investigation, L.S.P., and R.C. Resources, L.J.B., K.W.M., and A.J.H. Data curation, L.S.P. and R.C.

Writing - original draft, L.S.P. and R.C.

Writing - review & editing, L.S.P., R.C., and A.J.H. Visualization, L.S.P., R.C., S.L.S., and A.J.H.

Supervision, L.S.P., K.W.M., and A.J.H.

Project administration, L.S.P., K.W.M., and A.J.H. Funding acquisition, L.J.B., K.W.M., and A.J.H.

## Data and code availability

Microscopy data reported in this paper will be shared by the lead contact upon request. Bulk RNA sequencing data files have been deposited in the Gene Expression Omnibus (GEO) and GEO numbers will be made accessible upon final publication. Any additional information required to reanalyze the data reported in this paper is available from the lead contact upon request.

## Methods

### Mouse experiments

#### Animal care and handling

All mouse experiments followed National Institutes of Health (NIH) guidelines and were approved by the Institutional Animal Care and Use Committee (IACUC) of the University of Pennsylvania (protocol #800700). Embryos were collected from timed pregnant outbred CD-1 mice (Charles River Laboratories) that were housed in conventional facilities, kept on a standard diet and 12 hr light/dark cycle, and euthanized by CO_2_ inhalation. Embryos were dissected at embryonic day (E)12-15 in chilled Dulbecco’s phosphate buffered saline (1x DPBS, #MT21-31-CV, Corning) and dissected tissues were either processed immediately for immunofluorescence or cultured as explants. Embryo ages were roughly confirmed for each litter by limb staging ^67^.

#### Air-liquid interface culture

E13 kidney explants were cultured *ex vivo* on transparent 0.4 µm pore Transwell filter inserts (#3460, Corning) mounted in a 24-well plate ^45^. Explants were gently placed near the center of the insert with a cut P20 pipette tip and excess media was quickly removed to allow the explant to settle on the filter paper. The lower compartment of each well was filled with 250 µl live imaging medium, which consists of phenol red-free DMEM (4.5 g l^-1^ glucose, L-glutamine, and 25 mM HEPES, #21063-029, Invitrogen) supplemented with 10% FBS (#MT35-010-CV, Corning, Lot# 19321001), 1 mM sodium pyruvate (100 mM stock, #11360070, Invitrogen), and 100 U ml^-1^ penicillin-streptomycin antibiotic (1x pen-strep, 10,000 U ml^-1^ stock, #14150122, Invitrogen). Treatments were either 100 µg ml^-1^ recombinant human GDNF (#212-GD-050, R&D Systems) or 100 nM Selpercatinib (RETi). To visualize the ureteric bud, kidneys were pre-labeled with FITC-labeled anti-CD326 (EpCAM) antibodies (#11-5791-82, eBioscience) or eFluor-660 labeled anti-CD324 (ECAD) antibodies (#50-3249-80, eBioscience) diluted 1:250 in DMEM for 30-60 min and washed once in DMEM. Explants were cultured for 4 days with daily medium changes.

## Cell lines and organoids

### Human iPSC maintenance

Human iPSC studies were reviewed and approved by the University of Pennsylvania’s Institutional Biosafety Committee (IBC, protocol #26-074). Lines expressing endogenously tagged MAFB:TagBFP and GATA3:mCherry markers (iPSC^MAFB/GATA3^) are described previously ^57^ and were obtained from the Washington University Kidney Translational Research Center and Division of Nephrology. PENN123i-SV20 human iPSC line (SV20, male, WiCell, Lot # DB36624) was obtained from the Children’s Hospital of Philadelphia Stem Cell Core. All iPSCs were maintained in mTeSR™ Plus feeder-free medium (#100-0276, Stem Cell Technologies) with daily medium exchanges and were subcultured every 3-4 days using ReLeSR™ dissociation reagent (#100-0483, STEMCELL Technologies) for colony passaging, or Accutase (#07920, STEMCELL Technologies) for single cell passaging. All maintenance lines were cultured in a humidified incubator at 37°C and 5% CO_2_ in 6-well plates pre-coated with hESC-certified Matrigel (1:100 in DMEM, #354277, Corning). Genomic integrity for all cell lines was verified in a previous publication ^34^.

### iUB organoid differentiation

Human iPSCs were differentiated into iUB organoids over a 7-day process ^68^. First, a single cell suspension of 3.0-3.25x10^4^ cells was plated in each well of an hESC-Matrigel coated 24 well plate in 500 µl maintenance medium supplemented with 10 µM Y-27632 (ROCKi, #72304, STEMCELL). After ∼1 day, these were differentiated into primitive streak mesendodermal progenitors in a basal medium consisting of Advanced RPMI1640 (#12-633-012, Fisher) supplemented with GlutaMAX™ (#35050061, Invitrogen) supplemented with 50 ng ml^-1^ activin A (#338-AC, R&D Systems), 25 ng ml^-1^ bone morphogenic protein 4 (BMP4, #120-05ET, PeproTech), 5 µM CHIR99021 (CHIR, #13122, Cayman Chemical), and 25 ng ml^-1^ FGF2 (#100-18B, PeproTech) over a period of ∼25-27 hr ^69^. Over days 1-3, cells were differentiated into PIM in basal medium supplemented with 25 ng ml^-1^ FGF2, 1 µM A83-01 (TGFβi, #9001799, Cayman Chemical), 0.1 µM LDN193198 dihydrochloride (ALKi, #19396, Cayman Chemical), and 0.1 µM retinoic acid (RA, #R2625, Sigma). We routinely cryopreserved day 3 PIM cells ^35^ by lifting cells with Accutase, centrifuging, and freezing 1.5-3x10^6^ cells in freezing medium (45% basal medium, 45% FBS, 10% DMSO). Thawed aliquots resumed differentiation into iUB using the day of thawing as day 3. All culture and differentiation media components are specified in **Table S1**.

ND spheroids were cultured over days 3-7 in a 24-well microwell plate (AggreWell^TM^ 400 & AggreWell^TM^ 800, #34415 & #34815, STEMCELL). Prior to plating cells, microwell surfaces were passivated with Anti-Adherence Rinsing Solution (#07010, STEMCELL), centrifuged at 1,300 xg for 5 min, and washed with DMEM. Day 3 PIM cells were then resuspended in basal medium supplemented with 50 ng ml^-1^ FGF9 (#273-F9, R&D Systems) and 0.1 µM RA, and ∼5-8x10^5^ cells were seeded into each well and centrifuged at 100 xg for 3 min to form spheroids. On day 5, we performed a half medium exchange and cultured cells in basal medium supplemented with 50 ng ml^-1^ GDNF (#450-10, PeproTech) and 0.1 µM RA until day 7. Reagents and recombinant proteins used for human iPSC differentiation were handled and stored as described previously ^69^.

To embed iUB organoids, we pre-coated the bottom surface of 24-well plates (Nunclon Delta, #142475, Thermo Scientific) with ∼15 µl of growth factor reduced Matrigel matrix (GFR Matrigel, #354230, Corning) and allowed it to solidify at 37°C for >15 min before embedding. On day 7, spheroids were vigorously dislodged from microwells by pipetting with a P1000 tip for ∼30 sec, transferred to a 1.5 ml microcentrifuge tube, and allowed to settle by gravity for ∼5 min. We carefully aspirated excess medium resuspended spheroids in ∼650 µl of cold GFR Matrigel. A 45 µL droplet containing Matrigel and spheroids was deposited onto the Matrigel-coated surface using a wide-bore P200 pipette tip. For black walled 96-well plates (µCLEAR, #655090, Greiner) or 10-well black walled chambered slides (CELLview™, #53079, Greiner, Greiner) we used 7 µl of GFR Matrigel for the base layer and a 25 µl spheroid droplet volume. Plates were then incubated at 37°C for 30-60 min to allow Matrigel to solidify before adding medium.

Ureteric bud medium (UBM) used on days 7+ consists of basal medium supplemented with 2 µM CHIR99021, 50 ng ml^-1^ fibroblast growth factor 10 (FGF10), 1 µM A83-01, 0.1 µM LDN193187, 0.1 µM RA, 10 µM Y-27632, and 50 ng ml^-1^ GDNF. This formulation was used as the standard branching reference condition (+GDNF, ref. ^30^). The negative control medium (-GDNF) consisted of UBM without GDNF. The +RETi condition consisted of complete UBM supplemented with 100 nM Selpercatinib and the excess GDNF (++GDNF) condition contained 250 ng ml^-1^ GDNF. UBM and other media components are described in **Table S1**.

### Cell culture

Madin-Darby canine kidney epithelial cells (MDCK II, female, #00062107-1VL, Millipore Sigma) were maintained in minimum essential medium (MEM, Earle’s salts and L-glutamine, #MT10-010-CM, Corning) supplemented with 10% FBS (#MT35-010-CV, Corning, Lot# 19321001) and 100 U ml^-1^ penicillin-streptomycin (10,000 U ml^-1^ stock, #14150122, Invitrogen), and were passaged every 3-4 days using 0.25% trypsin-EDTA (#25200056, Corning). MDCK-RET and MDCK-KM cell lines (ref. ^54^) were routinely cultured with 100 µg ml^-1^ neomycin (G418, 50 mg ml^-1^ stock, #61-234-RG, Corning) to remove non-expressing cells.

Human embryonic kidney cells (HEK 293T, #632180, Takara Bio) were maintained in DMEM (4.5 g L^-1^ glucose, L-glutamine, and sodium pyruvate, #MT10-013-CV, Corning) supplemented with 10% FBS and 100 U ml^-1^ penicillin-streptomycin, and were passaged every 3-4 days using 0.05% trypsin-EDTA (#25200054, Corning). Live imaging medium consisted of phenol red-free DMEM (4.5 g l^-1^ glucose, L-glutamine, and 25 mM HEPES, #21063-029, Invitrogen) supplemented with 10% FBS, 1 mM sodium pyruvate (100 mM stock, #11360070, Invitrogen), and 100 U ml^-1^ penicillin-streptomycin. All cell lines were subcultured in polystyrene flasks and maintained in a humidified incubator at 37°C and 5% CO_2_. All culture plates and flasks were handled under red light and wrapped in aluminum foil to minimize blue light exposure.

## Plasmids and cloning

### Plasmids

Full length human RET and GFRA1 expression vectors are previously described ^70,71^ and include pcDNA3.1(+)-RET9-FL, pcDNA3.1(+)-RET51-FL, and CH269_GFRA1. Actin cytoskeleton dynamics were visualized by Lifeact fused to enhanced green fluorescent protein (eGFP) via pTK92_Lifeact-GFP (#46356, Addgene, RRID:Addgene_46356) or pTK93_Lifeact-mCherrry (#46356, Addgene, RRID:Addgene_46356) (ref. ^72^). Nuclei were visualized with histone 2B (H2B) constructs fused to a monomeric red fluorescent protein (mRuby2) via pLentiPGK Hygro DEST H2B-mRuby2 (#90236, Addgene, RRID:Addgene_90236) (ref. ^73^) or yellow fluorescent protein (mVenus) via pLentiPGK Hygro DEST H2B-mVenus (this paper). The pLentiPGK Hygro DEST H2B-mVenus vector was generated by replacing the mRuby2 coding sequence with mVenus using standard molecular cloning techniques (see: **Cloning**) and a full sequence is provided (see **Data and code availability**). Viral packaging and envelope plasmids included pCMV-dR8.91 (#12263, Addgene, RRID:Addgene_12263), pMD2.G (#12259, Addgene, RRID:Addgene_12259), pCMV-VSV-G (#8454, Addgene, RRID:Addgene_8454), and pCMV-gag/pol (#14887, Addgene, RRID:Addgene_14887). For piggyBac experiments, we used the Super PiggyBac transposase (#PB210PA-1, System Biosciences) and custom transposon vectors containing the optoRET sequence (see: **OptoRET design**).

### Cloning

Linear DNA fragments were produced by polymerase chain reaction (PCR) using Q5 polymerase (#M0492S, New England Biolabs) with cycle times and temperatures adjusted according to manufacturer instructions. PCR products were subsequently digested with DpnI (#R0176S, New England Biolabs) to remove template DNA and purified using a QIAquick PCR purification kit (#28104, Qiagen). pHR_CMV lentiviral vector was obtained by double restriction digest using MluI-HF® (#R3198S, New England Biolabs) and NotI-HF® (#R3189S, New England Biolabs). CLPIT retroviral vector was obtained by double restriction digest with NotI-HF® and SfiI (#R0123S, New England Biolabs). Vectors were purified by gel electrophoresis and extracted using a Zymoclean Gel DNA Recovery kit (#D4001, Zymo Research). All final constructs were assembled with NEBuilder® HiFi DNA Assembly Master Mix (#E2621, New England Biolabs), transformed into NEB® Turbo Competent *E. coli* (#C2984H, New England Biolabs), and purified with the ZR Plasmid Miniprep Classic kit (#D4015, Zymo Research).

### OptoRET design

We generated the optoRET tool by modifying an existing optogenetic RET expression vector (opto-hRET_317, #58747, Addgene, RRID:Addgene_58747) (ref. ^11^).

Opto-hRET_317 contains an N-terminal myristoylation (Myr) peptide for plasma membrane insertion followed by a 415 amino acid region of the human RET9 intracellular domain (ICD), the *Vaucheria frigida* AUREOCHROME1 LOV domain (VfAU1-LOV), and a C-terminal influenza haemagglutinin (HA) tag. We amplified a 434 aa region containing the Myr domain, a 5 aa linker sequence, and the entire RET(ICD) sequence by PCR and appended a monomeric red fluorescent protein (mCherry) and the *Arabidopsis thaliana* cryptochrome 2 photolyase homology domain (CRY2^PHR^) from a previously described optogenetic fibroblast growth factor receptor tool (optoFGFR, ref. ^74^) to obtain a 1171 aa sequence (Myr-RET(ICD)-mCherry-CRY2^PHR^), hereafter referred to as optoRET. We assembled optoRET into the pHR vector with a cytomegalovirus (CMV) promoter ^75^ or the CLPIT retroviral vector under control of a tetracycline-repressive (Tet-OFF) promoter ^51^. We also produced variants where we replaced mCherry with an eGFP coding sequence for spectral compatibility with other fluorophores. For piggyBac experiments, we ordered the optoRET transposon construct (pPB[Exp]-CAG>Myr-RET(ICD)-eGFP-CRY2^PHR^) as a custom vector (VectorBuilder Inc.). All optoRET sequences and primers are provided (see **Data and code availability**).

## Transfection and stable cell lines

### Transient transfection

HEK 293T cells were transiently transfected using FuGene HD transfection reagent (#E2311, Promega). One day before transfection, ∼10^5^ cells were plated in antibiotic-free culture medium in each well of a 24 well plate. Next day, cells at ∼70% confluence were transiently transfected by mixing 1 µg of total plasmid DNA with 3 µl FuGene HD in Opti-MEM (#31985062, ThermoFisher) and incubated 10 min before adding to the wells. We selected stable HEK 293T cell lines expressing GFRA1, RET9, and RET51 using 200 µg ml^-1^ hygromycin B (50 mg ml^-1^ stock, #10843555001, Roche) and 400 µg ml^-1^ neomycin/G418, replacing the antibiotic-containing medium every 2 days for ∼1 week. Stable lines were expanded and are characterized for RET and GFRA1 co-expression in **Fig. S7**.

### Viral transduction

To produce viral particles, we seeded 7x10^5^ packaging cells (Lenti-X™ 293T, #632180, Takara Bio) per well in a 6-well plate and transfected them with viral envelope, packaging, and transfer plasmids ∼1 day later. For lentiviruses, we co-transfected 1.5 µg pHR or pLenti transfer plasmid, 1.3 µg pCMV-dR8.91 and 0.17 µg pMD2.G in each well. For retroviruses we co-transfected 1.25 µg CLPIT or pBabe transfer plasmid, 0.75 µg pCMV-VSV-G, and 0.5 µg pCMV-gag/pol (ref. ^74^). Plasmid mixtures were diluted in DI water and buffered to 1x HEPES-buffered saline (HeBS, 2x stock: 50 mM HEPES, 280 mM NaCl, 1.5 mM Na_2_HPO_4_, pH 7.05) to a final volume of 300 µl, followed by dropwise addition of 18 µl of 2.5 M calcium chloride (CaCl_2_) to a final concentration of 150 mM. This mixture was incubated at room temperature for ∼1 min 45 sec before adding to the packaging cell medium. Fresh media was exchanged after 24 hr. After another 24-48 hr, the supernatant was collected, centrifuged at 800 xg for 3 min, and passed through a 0.45 µm syringe filter before immediate use or stored at -80°C.

To generate stable cell lines, 1-2x10^5^ target cells were seeded in each well of a 6-well plate along with 0.2-1 ml viral supernatant in medium supplemented with 8 µg ml^-1^ polybrene (#TR-1003-G, EMD Millipore). Unused viral aliquots were stored for up to 1 year at -80°C and were quickly thawed at 37°C before use. Stable cell lines were expanded and enriched for >90% expressing cells by fluorescence activated cell sorting (FACS) using either a BD Influx™ cell sorter (BD Biosciences) or a BD FACSAria™ III cell sorter (BD Biosciences). Cell lines expressing Lifeact-GFP were further selected for 3-4 days with 2.5 µg ml^-1^ puromycin dihydrochloride (10 mg ml^-1^ stock, #A11138-03, Life Technologies). Lines expressing H2B-mRuby2 or H2B-mVenus were further selected for ∼1 week with 200 µg ml^-1^ hygromycin B (50 mg ml^-1^ stock, #10843555001, Roche) to remove non-expressing cells.

### PiggyBac transfection

Human iPSC cell lines stably expressing optoRET (iPSC-optoRET) were generated using the piggyBac transposon system. First, cells were transiently transfected with the Super PiggyBac transposase and pPB[Exp] transposon vectors containing the optoRET sequence (see **Data and code availability**). One day before transfection, cells were seeded at a density of 5x10^4^ cells per well in a 24-well plate that was precoated with hESC-grade Matrigel (see: **human iPSC maintenance**). Next day, 1 µl of Lipofectamine™ Stem Transfection Reagent (#STEM00003, ThermoFisher) was mixed with 500 ng of total DNA (1:3 Super PiggyBac:pPB[Exp] ratio) in 50µl of Opti-MEM™ reduced serum medium (#31985062, ThermoFisher). Plasmid DNA:Lipofectamine complexes were incubated for 10 min at room temperature before adding to the cell culture medium and removed after overnight incubation.

Transfected cells were enriched for >90% optoRET+ cells using a BD FACSAria™ III cell sorter (BD Biosciences) and expanded into a stable iPSC-optoRET pool.

## RNA transcriptomics

### Xenium spatial sequencing

Publicly available Xenium spatial transcriptomics datasets of mouse embryonic kidney (E17) and human fetal kidney (week 20) were obtained from a previous study ^17^. Processed output files were opened and visualized using Xenium Explorer (v4.0, 10x Genomics). For all datasets, nuclei were visualized using the DAPI channel with intensity values displayed in grayscale and scaled from 0 to 10,000. Cell boundaries provided by the Xenium segmentation pipeline were used for visualization and analysis. Transcript molecules were displayed as individual point features corresponding to their spatial coordinates. Cell group annotations were generated within Xenium Explorer using k-means clustering (k = 3) based on cellular gene expression profiles. All visualization parameters were held constant across samples to enable qualitative comparison between mouse and human datasets.

### RNA-seq

Day 12 iUB or day 11 iUB-optoRET organoids were extracted from Matrigel droplets using 600 µl of chilled Cell Recovery Solution (#354253, Corning) on ice for 20-30 min and were triturated approximately every 5 min. Organoids from each group were pooled into a single 15 ml conical tube, washed briefly in 1x DPBS, and placed in DNA/RNA shield (#R1100, Zymo). All sample preparation areas were pre-cleaned with RNaseZAP (#R2020, Sigma) before handling. RNA concentration measurement, cDNA library preparation, and mRNA poly-A capture 3’ bulk sequencing was performed by Illumina NovaSeq sequencing using a commercial service (Plasmidsaurus) and samples were shipped according to their shipping instructions. Samples were otherwise stored at -80°C.

### Differential Gene Expression analysis

Bulk RNA-seq data were processed and analyzed as described previously ^17^. FASTQ file quality was assessed using FastQC (v0.12.1). Reads were then quality filtered using fastp v0.24.0 with poly-X tail trimming, 3’ quality-based tail trimming, a minimum Phred quality score of 15, and a minimum length requirement of 50 base pairs (bp).

Quality-filtered reads were aligned to the reference genome using STAR aligner (v2.7.11, Alexander Dobin) with non-canonical splice junction removal and output of unmapped reads, followed by coordinate sorting using samtools (v1.22.1). PCR and optical duplicates were removed using Unique Molecular Identifiers (UMI)-based deduplication with UMIcollapse (v1.1.0). Alignment quality metrics, strand specificity, and read distribution across genomic features were assessed using RSeQC (v5.0.4) and Qualimap (v2.3), with results aggregated into a comprehensive quality control report using MultiQC (v1.32) (ref. ^76^). Gene-level expression quantification was performed using featureCounts (subread package v2.1.1) with strand-specific counting, multi-mapping read fractional assignment, exons and 3’-UTR as the feature identifiers, and grouped by *gene_id*. Final gene counts were annotated with gene biotype and other metadata extracted from the reference GTF file. Differential expression analysis was performed with edgeR (v4.0.16) (ref. ^77^) using a paired analysis by experimental replicate, including filtering for low-expressed genes with *edgeR::filterByExprwith* default values. Density plots along the log_2_(Fold Change) axis were determined for the top 7000 variable genes using kernel density estimation. Gene set enrichment analysis was performed by pulling gene sets from Hallmark, Reactome, and Gene Ontology databases from the human Molecular Signature Database (MSigDb) (refs. ^78,79^).

## GDNF and optogenetic stimulation assays

### GDNF stimulation

Black walled 96 well plates (µCLEAR, # 655090, Greiner) were pre-coated with 10 µg ml^-1^ bovine serum fibronectin (#F1141, Sigma) diluted in 1x DPBS for ∼20-30 min.

For HEK 293T experiments (**Figs. S7,8**) we plated 15,000 cells per well. For MDCK experiments (**Fig. 3** and **Fig. S9,10**) we plated 5,000 cells per well. Plates were centrifuged for 1 min at 20 xg to facilitate even distribution across the bottom and cells were allowed to adhere overnight. Next day, HEK 293T cells were serum starved for 5-6 hr before treatment with 100 ng ml^-1^ GDNF or medium (control) for 2 hr. MDCK cells were not starved and were treated with 50 ng ml^-1^ GDNF (#212-GD-050, R&D Systems) and 100 ng ml^-1^ recombinant human Gfrɑ1 (#714-GR-100, R&D Systems) at defined time points. Cells were kept in a humidified incubator at 37°C and 5%CO_2_ for the duration of stimulation.

### OptoPlate-96 stimulation

15,000 HEK cells or 2,000 MDCK cells were plated in black-walled 96-well plates (µCLEAR, #655090, Greiner) as described above. MDCK cells (**Fig. 3** and **Fig. S9,10**) were partially serum starved in media containing 0.5% FBS for 10-12 hr prior to stimulation. MDCK cysts were plated as described previously (see: **MDCK cysts** and **collagen I gels**). Inhibitors (100 nM Trametinib or 100 nM Selpercatinib) or DMSO (vehicle) were added to wells immediately prior to stimulation. For iUB-optoRET 2D culture experiments (**Fig. 4B-D**), spheroids were resuspended in UBM under the indicated conditions and distributed into 96-well plates. Plates were centrifuged at 100 × g for 1 min, incubated overnight to allow spheroid spreading, and subsequently stimulated. For 3D iUB branching experiments, wells in black-walled 24-well plates (#1.5H coverslip, P24-1.5H-N, Cellvis) were pre-coated with 10 µL GFR Matrigel and allowed to solidify at 37°C. Spheroids were resuspended in cold GFR Matrigel, dispensed as droplets positioned directly above the underlying optoPlate-96, and incubated for 30–60 min at 37°C before adding UBM. During all steps, cells and organoids were handled in red light conditions and plates were wrapped in aluminum foil during incubation and covered by a lid during stimulation to minimize light exposure.

Blue light stimulation was carried out in the optoPlate-96 using an array of 470 nm LEDs (#LBT64GV1CA59Z or #LB T64G-AACB-59-Z484-20-R33-Z, ams OSRAM AG) with individual stimulation instructions provided by a programmable Arduino Micro microcontroller (#A000053, Arduino) as described previously ^52^. We used two different optoPlate-96 formats: a 1-color blue version (max ∼350 mW cm^-2^) or a 3-color version (max ∼60 mW cm^-2^). For each, we programmed the light intensity, stimulation duration, cycle times, and LED intensities (reported in figure legends) using Arduino IDE software (v2.3.2). Individual 470 nm LEDs were calibrated for each plate using a handheld light meter (PM16-140 Thorlabs). All experiments were carried out in a humidified incubator at 37°C and 5% CO_2_.

### Whole field laser stimulation

MDCK cells used in optogenetic scattering experiments (**Fig. 3G,H** and **Fig. S10**) were imaged on a spinning disk confocal microscope using the 488 nm laser line for stimulation and 514/594 nm lines for imaging mVenus and mCherry fluorophores, respectively (see: **Spinning disk confocal microscopy**). Images were acquired in the 514/594 nm channels at a single plane across 2-3 positions per well every 5 min for ∼12 hr. We used the ND Sequence Acquisition function to separate the *xy* coordinates of regions receiving 488 nm stimulation (+light) from control regions (-light). Each *xy* position in the +light group received a single 488 nm laser pulse at 20% power (2.16 mW, 1 s duration). For imaging H2B-mVenus we used the 514 nm line at 5% power and 100 ms exposure and measured a maximum 475 nm output of 0.35 mW. We calibrated each line at 475 nm at the focal plane of a 20x lens using a handheld light meter (PM100D & S120VC, ThorLabs).

### DMD stimulation

We stimulated iUB-optoRET organoids using a DMD-equipped spinning disk confocal microscope (see: **Spinning disk confocal microscopy**). DMD stimulation experiments were carried out in NIS Elements software (Nikon, v5.11.00) JOBS using a dedicated script. First, for each experiment, *xyz* coordinates were identified for individual organoids within an experiment. These were used to capture a 170 µm z-stack (30 µm slices) of each organoid every 2 hr. After all z-stacks were acquired, an internal loop stimulated a subset of these *xyz* positions in a single plane using the DMD within a user-defined ROI. This internal stimulation loop (488 nm pulses, 5% power, 0.5 s exposure) was carried out at 4 min intervals thirty times per cycle. The script concatenated each *xy* position into a single time series stack at the end of the experiment, which were used for further analysis.

### DMD/optoPlate-96 Calibration

To calibrate DMD illumination intensity relative to the optoPlate-96, 10 µm diameter fluorescein isothiocyanate (FITC)-labeled polystyrene microspheres (#PS10u-FC-1, Nanocs) were mixed in a suspension containing 70% Matrigel, 20% DMEM, and 10% bead solution and plated in black-walled 96-well plates (µCLEAR, #655090, Greiner). 30 µl droplets of bead mixture were deposited in each well and allowed to gel for 30 min, after which 150 µl DPBS was added to cover the beads.

Photobleaching experiments were performed on a spinning disk confocal microscope equipped with a digital micromirror device (DMD) using a 10x objective. Experiments were carried out in NIS Elements software (Nikon Instruments) using a dedicated JOBS script. First, xyz coordinates were identified for imaging positions containing embedded beads. The script then acquired fluorescence images in the 488 nm channel from all positions prior to stimulation (pre-exposure images). A predefined square DMD region of interest was used to deliver continuous 488 nm illumination at 5% laser power. The JOBS script sequentially stimulated specified imaging positions for defined durations (3, 6, 12, or 18 min), while control positions received no stimulation (0 min). After all stimulations were complete, the script acquired fluorescence images of the same positions using identical imaging settings (post-exposure images).

To compare DMD illumination with the optoPlate-96, bead-containing wells were prepared as described above. Wells were illuminated on an optoPlate-96 for 12 min at LED intensities of 2.5, 5, 10, 25, 50, 100, 200, 250, 290, or 320 mW cm^-2^. Fluorescence images were acquired before and after illumination using the same 488 nm imaging settings and analyzed using the same background-corrected fluorescence quantification described above.

## MDCK morphogenesis assays

### 2D scattering

2,000 MDCK cells were seeded in each well of a 96-well plate (µCLEAR, # 655090, Greiner) that had been pre-coated with 10 µg ml^-1^ fibronectin. Plates were centrifuged for 1 min at 20 xg to settle cells. Approximately 10-12 hr before the onset of imaging, the media was replaced with a low-serum (0.5% FBS) imaging medium. For ligand stimulation of MDCK-RET or MDCK-KM cells, we added medium containing 50 µg ml^-1^ GDNF and 100 µg ml^-1^ Gfrɑ1 approximately 1 hr prior to the onset of imaging. Control medium contained 100 µg ml^-1^ Gfrɑ1 alone. For inhibitor experiments with MDCK-optoRET cells we added 100 nM Trametinib (MEKi) or 100 nM Selpercatinib (RETi) at the onset of imaging.

### MDCK cysts

MDCK cysts were pre-formed in growth factor reduced Matrigel following an established protocol ^55^. First, 30 µl of chilled Matrigel was spread across each well of a 24-well polystyrene plate and transferred to 37°C for ∼30 min to solidify. MDCK cells were passed with 0.25% trypsin-EDTA, dissociated into a single cell suspension, and mixed 1:1 with Matrigel to a 400 µl final volume containing ∼10^4^ cells. 200 µl of the cell:Matrigel mixture was pipetted into each coated well and the plate was placed at 37°C for 20-30 min before filling the wells with 1 ml complete medium. Cysts were grown for 4-6 days, replacing fresh culture medium once.

### Collagen gels

We then transferred cysts to collagen gels in either 8-well chambered slides (#CCS-8, Mattek), 8-well µ-slides with a No. 1.5 uncoated polymer coverglass bottom (#80821, Ibidi), or black-walled 10-well chambered slides (CELLview™, #53079, Greiner). Rat tail type I collagen (#354226, Corning) was diluted to 2 mg ml^-1^ in sterile DI water buffered with 10x DPBS (10x stock, #1420075, Gibco) on ice and adjusted to pH 7.0 with 1 N NaOH. To visualize individual collagen fibers, we pre-labeled 1 ml of type I collagen with Alexa Fluor™ 647 N-hydroxysuccinimidyl (NHS) ester (1:1000, #A20006, ThermoFisher), incubated for >12 hr at 4°C, and mixed with unlabeled collagen (1:4 labeled:unlabeled ratio). Each well was pre-coated with ∼30 µl neutralized collagen and placed in a 37°C incubator for ∼10 min. To remove cysts from Matrigel, we aspirated the medium, vigorously washed each well with chilled DPBS++ (1xDPBS with Ca^2+^/Mg^2+^, #MT21-030-CM, Corning) totaling 5 ml per well, and centrifuged the entire solution at 500 xg for 3 min. The ensuing slurry of cysts and matrix was resuspended in 10 ml DPBS++, incubated in a bucket of salted ice for ∼30 min to dissolve Matrigel, centrifuged at 500 xg for 3 min, and resuspended into a ∼50 µl volume. Cysts were mixed with the collagen solution and 100 µl of the cyst:collagen mixture was added to each well. To stimulate branching, we added 100 µg ml^-1^ Gfrɑ1 and 50 µg ml^-1^ GDNF after ∼16-24 hr or used the optoPlate-96 (details provided in figure legends and in **OptoPlate-96 stimulation**).

## Imaging

### Spinning disk confocal microscopy

Live and fixed tissues were imaged on one of two spinning disk confocal systems built on a Ti2-E microscope stand (Nikon Instruments) and equipped with a spinning disk confocal scanhead (CSUW-1, Yokogawa). The first system was equipped with a motorized stage, perfect focus system (PFS) for stable z-plane localization, Prime BSI sCMOS camera (Teledyne Photometrics), high power white light LED for brightfield imaging, and laser illumination provided by 405, 488, 561, and 640 nm lines (100 mW each) passed through standard single band pass filter sets (#ET455/50m, #ET525/36m, #ET605/52m, and #ET705/72m, Chroma). Images were acquired through 4x/0.20 NA, 10x/0.45 NA, or 20x/0.75 NA Plan Apo λ objectives, or an S Plan Fluar ELWD 20x/0.45 NA objective (Nikon) under control of NIS Elements AR software (Nikon, v5.11.00). A separate light path contained a digital micromirror device (DMD, Polygon 400, Mightex), multi-wavelength LED light source (Spectra III, Lumencor) and a C-TIRF Ultra High Signal-to-Noise 488 nm Total Internal Reflection Fluorescence (TIRF) filter set (Chroma) to provide even 488 nm illumination at the focus plane. A custom-built environmental enclosure (OkoLab) maintained environmental control at 37°C and 5% CO_2_ for live sample imaging.

A second spinning disk system was also built on a Nikon Ti2-E stand with motorized stage, PFS, CSUW-1 scanhead (Yokogawa), ORCA-Fusion sCMOS camera (Hamamatsu), high power white light LED for brightfield imaging, and laser illumination provided by 405, 445, 488, 514, 561, 594, and 640 nm laser lines (100 mW each) passed through appropriate single pass filter sets (#ET455/50m, #ET485/25M, #ET525/36m, #ET540/30m, #ET605/52m, #ET625/30M, and #ET705/72m, Chroma) and separate dichroic mirrors for 405/488/561/640 nm and 445/514/594 nm lines. Images were acquired through 4x/0.20 NA, 10x/0.45 NA, and 20x/0.8 NA Plan Apo λD objectives or a S Plan Fluar LWD 20x/0.7 NA objectives (Nikon) under control of NIS Elements AR software (Nikon, v5.11.00). Samples were kept at 37°C and 5% CO_2_ in a humidified enclosure (OkoLab) for the duration of imaging.

### Epifluorescence microscopy

Fixed and live cells were imaged directly on culture plates using an epifluorescence microscope built on a Ti2 stand (Nikon) and equipped with a motorized stage, PFS, and high power white light LED for transmitted illumination. Epifluorescence illumination was provided by a Sola SEII 365 light engine (89 North) run through single bandpass filter sets (#DAPI C-FL, #GFP-4050-000, #TRITC-B-000, and #LED-Venus-A-ZERO, Semrock; #49022-ET-Cy5.5, Chroma) and 4x/0.20 NA, 10x/0.45 NA, and 20x 0.8NA Plan Apo λ objectives or an S Plan Fluar ELWD 20x/0.45 NA objective (Nikon) under control of NIS Elements AR software (Nikon, v5.11.00). Live samples were kept at 37°C and 5% CO_2_ in a humidified enclosure (OkoLab) for the duration of imaging.

## Immunofluorescence

### Whole mount kidneys

We immunostained embryonic kidney tissues using a previous protocol^80^. Briefly, freshly dissected or ALI cultured kidneys were fixed at room temperature in 4% paraformaldehyde (PFA, #15170, Electron Microscopy Sciences) for 15 min followed by three washes in 1x DPBS (5 min each) and blocked for at least two hr in a blocking buffer consisting of 1x DPBS containing 0.1% Triton X-100 (#T9284, Sigma) and 5% donkey serum (#D9663, Sigma). Fixed and blocked kidneys were incubated with primary antibodies for 2 days at 4°C and fluorescent secondary antibodies for 2 days at 4°C, with three washes in PBS-Tx (1x DPBS + 0.1% Triton X-100) totaling at least 16 hr following each incubation. Primary antibodies and dilutions are reported in **Table S2**, all secondary antibodies were raised in donkey and were used at 1:300 dilution. Whole mount kidneys were cleared in a solution of one part benzyl alcohol and benzyl benzoate (BABB) before imaging ^81^. BABB consists of one part benzyl alcohol to two parts benzyl benzoate (#AC105862500, ThermoFisher). Samples were first dehydrated in a series of 5 min incubation steps in increasing concentrations of methanol (25%, 50%, and 75% v/v diluted in PBS-Tx) before switching to a 100% methanol solution. Dehydrated samples were then moved to the microscope stage before aspirating methanol and adding a droplet of ∼100 µl of BABB solution. Samples were imaged directly in BABB and recovered by extraction in a reverse methanol series. Fixed explants on transwell filters were transferred to glass bottom dishes for imaging.

### 2D iPSC cultures

iPSCs or day 0-3 cells were fixed in 4% PFA diluted in PBS for 45 min, then washed three times in PBS for 5 min each. Cells were blocked for 1 hr at room temperature in PBS containing 0.1% Triton X-100 (PBS-Tx) and 5% donkey serum then incubated in primary antibodies diluted in blocking buffer overnight at 4°C and secondary antibodies for 1 hr at room temperature, with three PBS-T (PBS + 0.1% Tween 20) washes between each incubation step. Primary and secondary antibodies are reported in **Table S2**, secondary antibodies were all used at 1:500 dilution.

### 3D iUB organoids

Day 7-12 iUB organoids in Matrigel were incubated in 600 µl of chilled Cell Recovery Solution (#354253, Corning) on ice for 20-30 min. Organoids were released from Matrigel by gentle trituration with a wide bore P1000 tip, collected in 15 ml tubes that had been pre-coated with Anti-Adherence Rinsing Solution, and allowed to settle by gravity. Pooled organoids were briefly washed 1x DPBS and centrifuged again, then fixed in 4% PFA diluted in PBS for 30-60 min at room temperature. Fixed organoids were transferred to a 24 well plate, washed twice in PBS-glycine (100 mM, #G8898, Sigma), and twice in PBS for 5 min each, blocked for 2 hr at room temperature, then transferred to a 96-well plate. Organoids were incubated in primary antibodies overnight at 4°C and secondary antibodies for 2 hr at room temperature with each step followed by three washes in PBS-Tx for at least 10 min each.

Blocking and antibody dilution buffer consisted of PBS-Tx with 5% donkey serum. Primary and secondary antibodies are reported in **Table S2**, secondary antibodies were all used at 1:500 dilution.

### 2D quantitative ppERK immunofluorescence

Cells were fixed in 4% PFA by adding 16% PFA directly to cell culture medium and incubating for 10 min at room temperature immediately following stimulation. Fixed cells were permeabilized in 1X DPBS containing 0.5% v/v Triton X-100 for 10 min at room temperature, and post-fixed in pre-chilled methanol for 10 min at -20°C. We omitted the methanol post-fixation step for MDCK experiments involving phalloidin staining (**Fig. 3D-E**). Samples were blocked in 1x DPBS containing 0.1% v/v Tween-20, (#9480, EMD Millipore) and 1% w/v BSA (#A2153, Sigma) for 30-60 min at room temperature, then incubated in primary antibodies overnight at 4°C and secondary antibodies for 1 hr at room temperature with 5 PBST washes following each incubation step. Secondary antibodies were diluted 1:1000 in blocking buffer and 300 nM DAPI (4′,6-diamidino-2-phenylindole dihydrochloride, #D1306, ThermoFisher) was added to the secondary antibodies to counterstain nuclei. Primary and secondary antibodies and dilutions are reported in **Table S2**.

### 3D MDCK cyst immunofluorescence

Cysts in collagen I gels were fixed in 2% PFA for 20 mins at room temperature by adding 16% PFA directly to the culture medium. Fixed tissues were washed three times with PBS containing 100 mM glycine (#G8898-500G, Sigma) and twice in PBS (5 min each wash), permeabilized in PBS with 0.5% Triton X-100 for 15 min, and blocked for 2 hr at room temperature in blocking buffer (PBS, 1% w/v BSA, and 0.1% v/v Triton X-100) supplemented with 5% donkey serum. Primary and secondary antibody incubations were each carried out overnight at 4°C with three 1 hr washes in PBS-Tx between each step. Primary and secondary antibodies are reported in **Table S2**. Secondary antibodies (all raised in donkey) were added at 1:500 dilution. To visualize nuclei and F-actin, we incubated with 300 nM DAPI and Alexa Fluor™ 488 phalloidin (1:100, #A12378, Invitrogen) for at least 1 hr before imaging.

## Image Analysis

### Kidney explant morphology

All images were analyzed using Fiji/ImageJ2 ^82^ (version 2.16.0) unless otherwise noted. Total kidney areas were manually traced from 4x brightfield images using the *Polygon Sections* tool, excluding the ureter (**Fig. S2A,B**). Epithelial structures were identified as EpCAM+ terminal branches on day 1 and RET+/EpCAM+ terminal branches on day 4 and used to annotate tips (**Fig. S2C**). In the +RETi samples, which lack tip cells, we identified tips using EpCAM/ECAD fluorescence and morphology. To measure RET+ tip cluster size (**Fig. S2D-F**), we first subtracted a fixed background value and used the *Filter>Median* tool (2x2 pixel kernel) to remove noise. We then thresholded the resulting image to create a binary mask of RET signal and used the *Analyze Particles* function with a minimum size cutoff of 260 µm^2^ (100 pixels^2^) to isolate features and manually curated images to remove non-physiological structures.

To measure ppERK and RET, we generated z-stacks at 20x magnification (2.5 µm per slice) by spinning disk confocal microscopy.We selected medial sections where lumens were visible and used the *Polygon Sections* tool to manually trace ROIs around the outline of individual cells using ECAD signal. Mean ppERK and RET signal and area of each ROI were extracted using the *Multi-Measure* tool. For +RETi samples that lacked a distinctive tip population, we defined the tip as the extent of ECAD+ cells surrounded by SIX2+ cap domain (**Fig. S5A**).

### Organoid morphology

Organoid morphology was processed from 10x stacks (5 µm per slice) acquired on the spinning disk or epifluorescence microscopes. To measure projected organoid area, we generated maximum intensity projections from these stacks and used the *Polygon Sections* tool to manually outline an ROI around the epithelial (GATA3+) organoid compartment boundary, then used the *Multi-Measure* tool to extract total area (µm^2^) and circularity of each region (**Fig. 1K,L**; **Fig. S4G,H**). Circularity is calculated as:

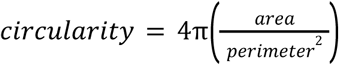

We quantified tip number by manually counting bud-like structures from either brightfield or mCherry maximum intensity projections.

### MDCK cyst morphology

We manually drew ROIs around individual cysts using the F-actin (phalloidin) channel with the *Polygon Sections* tool and calculated the mean optoRET (mCherry) fluorescence as well as the area and circularity of each cyst using the *Multi-Measure* tool. Cysts were excluded if they had clearly attached to and spread on the dish bottom or when multiple cysts could not be unambiguously identified from their neighbors.

### Single-cell ppERK quantification

We quantified fluorescence intensities from 3-5 fields per well and used a custom CellProfiler script ^83^ to measure fluorescence intensity values (ppERK, RET, GFRA1, and/or optoRET) for individual cells. Briefly, the script identifies nuclei from the DAPI counterstained image using the *IdentifyPrimaryObjects* module and Otsu thresholding. Next, the script dilates these objects to produce a 5-pixel-wide region of interest (ROI) around each nucleus using *IdentifySecondaryObjects* and *IdentifyTertiaryObjects* functions. We then subtract the background (lower quartile intensity) from each image as measured using the *ImageMath* function. The average grayscale intensity for each ROI is measured (*MeasureObjectIntensity*) from this background subtracted image. Data were subsequently filtered using R (v4.2.0, R Core Team) running on RStudio (v2023.06.0, Posit Software) to remove all objects where the fluorescence intensity of any channel was zero, since these typically represent debris. Image data was then rescaled to 16-bit depth and we measured the mean (*µ*) and standard deviation (*σ*) RET or optoRET fluorescence. Cells were counted as positive if the fluorescence signal of RET (*I_RET_*) or optoRET (*I_optoRET_*) was higher than the mean plus two standard deviations (*I_RET_>µ_RET_+2σ_RET_*). We directly compared ppErk levels for all RET+ or optoRET+ cells within an experiment across all treatment groups.

To estimate the optoRET tool sensitivity as a function of 470 nm intensity (**Fig. S8G,H**), we fitted the mean *ppERK* signal of each experiment to a Hill function:

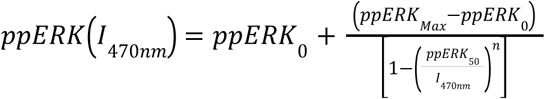

The minimum measured intensity (*ppERK_0_*), the fluorescence intensity at saturation (*ppERK_Max_*), the half-saturation value (*ppERK_50_*), and the Hill coefficient (*n*) are calculated as independent fitting parameters. We used R (version 4.20.0, R Core Team) running on RStudio (version 2023.06.0, Posit Software) to fit the measured mean *ppERK* signal of three replicate wells per condition for two independent experiments using the nonlinear least squares regression (*nls*) function. We estimated *ppERK_50_* occurs between 3.8 ± 0.25 and 7.6 ± 0.9 mW cm^-2^ [mean ± s.e.m.]. Model-predicted values for *ppERK_0_*(experiment 1: 219±143 a.u.; experiment 2: 328 ± 91 a.u., mean ± s.e.m.) were not significantly different from measured ppErk intensities for un-stimulated cells (experiment 1: 290 ± 506 a.u.; experiment 2: 278 ± 459 a.u., mean ± s.d.).

### Cell tracking

SIngle-nuclei tracking of scattering MDCK cells (**Fig. 3G,H, Fig. S9E-G**, and **Fig. S10**) was performed using ImageJ/Fiji using the TrackMate plugin ^56,84^ using the StarDist detection algorithm ^85^ for segmentation. We pre-processed images by isolating the nuclear (H2B) fluorescence channel from time-lapse stacks and used the *Filter>Median* (2x2 kernel) function to smoothen images. Individual nuclei tracks were extracted using a Linear Assignment Problem (LAP) tracker algorithm ^86^ with a maximum frame-to-frame distance of 30 µm.

Experiments were performed in two biological replicates with 2 wells (technical replicates) per condition.Nucleus *x,y* coordinates for each cell were used to calculate mean cell speeds and converted to relevant units (µm min^-1^). Each dataset was filtered using a custom code written in R (v4.20.0, R Core Team) running on RStudio (v2023.06.0, Posit Software) to remove tracks with fewer than 6 hr of consecutive timepoints.

### iUB organoid average projections

Normalized average projections were produced from the GATA3:mCherry channel using *Filter>Gaussian Blur* (2x2 kernel), *Math>Subtract*, and *Normalize* (Clijx toolbox) functions in ImageJ/Fiji. Individual projections were then processed using a second average projection step and displayed using the mpl-magma lookup table.

Tubule orientation angles were manually annotated using the *line* tool based on lines drawn from the organoid center to points at the tubule tips. Radial histograms were generated in R using *ggplot2* by binning bud angles into 30° intervals and plotting their frequency in polar coordinates. Statistical analysis was performed using the non-parametric Kolmogorov-Smirnov test, which compares the empirical cumulative distribution function (CDF) of bud angles to a reference distribution (uniform distribution). A significant value (*p* < 0.05) here indicates the likelihood of failing to reject the null hypothesis that bud angles are uniformly distributed.

### Bead photobleaching quantification

Fluorescence intensities of FITC beads were quantified in ImageJ/Fiji. Five beads from each of two wells within the DMD ROI were randomly selected, and circular ROIs were drawn around each bead and nearby background regions before and after stimulation. Percent fluorescence remaining after stimulation was calculated as:

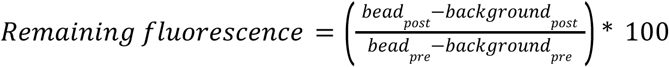

Percent fluorescence values were plotted as a function of exposure time in R (v4.2.0, R Core Team) running on RStudio (v2023.06.0, Posit Software) with the *ggplot2* package.

## Statistics and reproducibility

### Reproducibility and replication

All experiments were performed in biological and technical replicates, with the total number of individual measurements (n) and the number (N) and type of replicates reported in the figure legends. For experiments with mouse kidneys, distinct litters were counted as biological replicates while individual kidneys were counted as technical replicates. For cell or organoid experiments, individual passages or batches of differentiated cells were counted as biological replicates. Technical replicates within an experiment correspond to individual wells of a plate. Experiments were not randomized and investigators were not blinded to experimental conditions during assessment.

### Statistical analysis

All statistical analyses were performed using R (v4.2.0, R Core Team) running on RStudio (v2023.06.0, Posit Software). Summary plots and SuperPlots ^87^ were constructed in Tidyverse ^88^ using the *ggplot2* library. Before selecting an appropriate statistical test for each dataset, we first performed a Shapiro-Wilk normality test to assess whether the data were normally distributed (*p*<0.05 threshold). We used Welch’s two-sided t-test or one-way ANOVA for unpaired, normally distributed, independent samples with Tukey-Kramer *post hoc* test for multiple comparisons, or Dunnet’s test for multiple comparisons against a single reference group. In experiments where data were not normally distributed, we chose appropriate non-parametric tests such as the Kolmogorov-Smirnov test. For multiple groups of non-normally distributed data we used the Kruskal-Wallis test with Dunn’s *post hoc* test, using the Holm-Bonferroni method for multiple comparisons. Unless otherwise noted, our threshold for statistical significance was *p* < 0.05. Specific tests and *p*-values for each experiment are reported in figure legends.

