## Supplementary Information for "Synthetic budding morphogenesis by optogenetic receptor tyrosine kinase signaling"

### Supplementary Figures

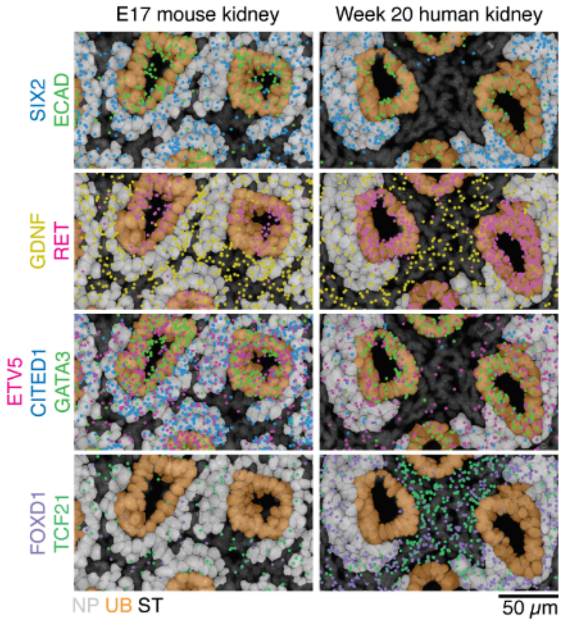


**Figure S1.** Xenium spatial sequencing of GDNF-RET signaling components in mouse and human kidney tissues.

*Left column*, Xenium spatial sequencing from a previously published dataset [17] of E17 mouse kidney showing localization of mRNA transcripts in nephron progenitor (NP), ureteric bud (UB), and stromal (ST) cells. *Right column*, spatial sequencing of week 20 human kidney from the same publication. Transcripts include epithelial cells (ECAD, GATA3), cap mesenchyme (SIX2, CITED1), ureteric bud tip (RET, ETV5), and stromal cells (FOXD1, TCF21).

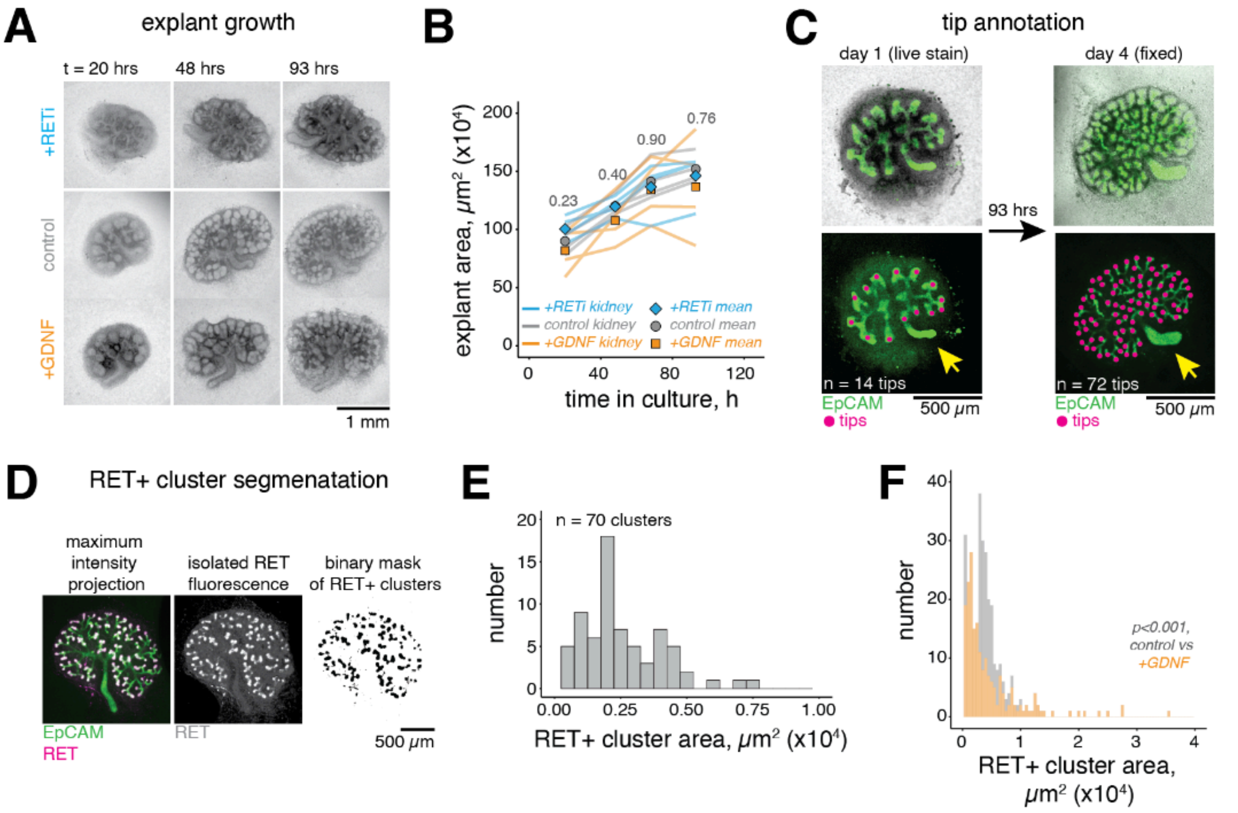


**Figure S2.** Annotations and quantification of kidney explant morphology.

A. Time series of E13 kidneys grown in ALI culture for 4 days (93 hrs) in the presence of 100 nM Selpercatinib (+RETi), 100 ng $ml^{-1}$ GDNF (+GDNF), or control media.

B. Explant areas ($\mu m^2$) measured on days 1-4 for the conditions above. Points are means of n = 4 explants per group at each timepoint, lines are individual explants. *P*-values by one-way ANOVA with Tukey's *post hoc* test, no significant comparisons ($p>0.05$) between conditions on any timepoint.

C. Live E13 kidney after 20 hrs culture on a transwell filter (day 1) and the same kidney after fixing and immunostaining at 93 hrs in culture (day 4). *Top row,* brightfield and epithelial structures labeled with a FITC-EpCAM antibody suitable for live imaging (ref. [45]). *Bottom row*, isolated FITC-EpCAM signal. Points are annotated tip locations, yellow arrow marks the ureter. Data are representative of measurements used to generate **Fig. 1F**.

D. Annotation of RET+ structures from an example E13 kidney grown in ALI culture and fixed on day 4. *Left*, maximum intensity projection of ECAD and RET signal following background subtraction. *Middle*, projected images were median filtered (2x2 kernel) to remove noise and isolate RET+ clusters. *Right*, binary masks were created by thresholding the RET signal to a fixed value and manually annotated to remove non-physiological structures.

E. Histogram of RET+ cluster area of n = 70 clusters for the kidney shown in panel d (mean ± s.d.: $0.26 \pm 0.16$ $(x10^4)$ $\mu m^2$).

F. Histogram of RET+ cluster area for n = 266,196 clusters (control, +GDNF) collected from N = 4 kidneys per condition. No clusters detected (n.d.) in +RETi kidneys. *P*-value by Kolmogorov-Smirnov test.

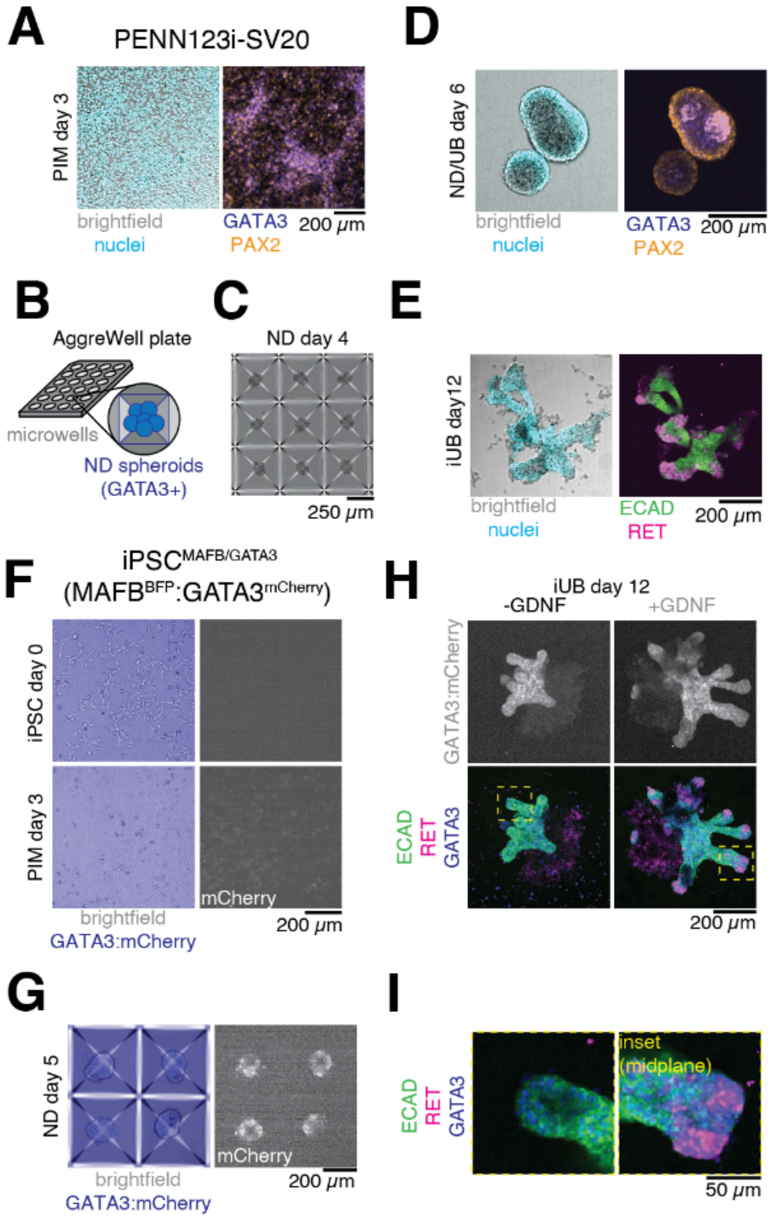


**Figure S3.** Differentiation of human iPSCs into iUB epithelial organoids.

A. PENN123i-SV20 (SV20) human iPSCs differentiated into day 3 PIM. *Left*, brightfield and nuclei (DAPI). *Right*, early kidney markers PAX2 and GATA3.
B. Multi-well plates (AggreWell) used to produce iND spheroids.
C. Day 4 ND spheroids produced from SV20 iPSCs in an AggreWell 400 plate.
D. Day 5 ND spheroids produced from SV20 iPSCs and removed from AggreWell plates. *Left*, brightfield and nuclei (DAPI). *Right*, immunofluorescence of early kidney markers PAX2 and GATA3.
E. Day 12 iUB organoids produced from SV20 iPSCs, extracted from Matrigel domes. *Left*, brightfield and nuclei (DAPI). *Right*, immunofluorescence of epithelial markers (ECAD) and tip markers (RET) showing tip-trunk hierarchy.

F. MAFB$^{BFP}$:GATA3$^{mCherry}$ dual reporter [57] iPSCs (iPSC$^{MAFB/GATA3}$) differentiated from day 0 into day 3 PIM. *Left*, live GATA3 reporter (mCherry) shown as an overlay with brightfield. *Right*, isolated greyscale images of mCherry.
G. Day 5 ND spheroids produced from iPSC$^{MAFB/GATA3}$ dual reporter iPSCs in AggreWell 400 plates. *Left*, live GATA3 reporter (mCherry) shown as an overlay with brightfield. *Right*, isolated greyscale images of mCherry.
H. Day 12 iUB organoids produced from iPSC$^{MAFB/GATA3}$ dual reporter iPSCs and cultured in UBM containing 0 ng $ml^{-1}$ GDNF (*left*, -GDNF) or 50 ng $ml^{-1}$ GDNF (*right*, +GDNF) from days 7-12. *Top*, maximum intensity projection of live GATA3 reporter. *Bottom*, immunofluorescence for epithelial junctions (ECAD), tip cells (RET), and endogenous GATA3. Insets of tip domains (*right*) show loss of RET+ tip cells in the -GDNF organoids and the presence of a robust tip domain in +GDNF ones.

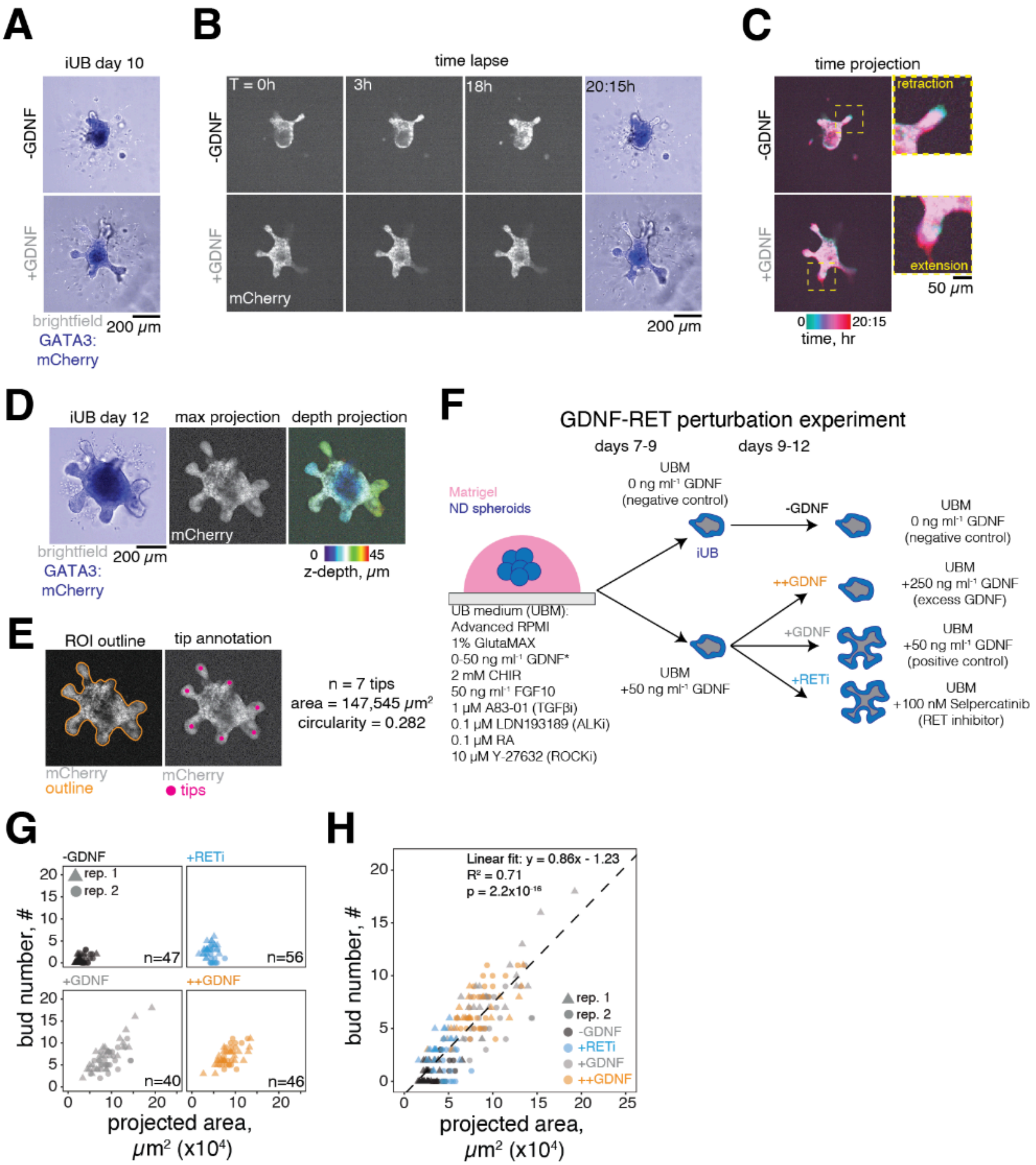


**Figure S4.** Morphology and budding quantification of iUB organoids

A. Day 10 iUB organoids grown from iPSC$^{MAFB/GATA3}$ cells. Organoids were grown in Matrigel domes in a 10-well chambered slide from day 7 in UBM with 0 ng ml$^{-1}$ GDNF (-GDNF) or 50 ng ml$^{-1}$ GDNF (+GDNF).
B. Time lapse images of GATA3 reporter (mCherry) from individual iUB organoids acquired at 45 min intervals over 20 hrs, 15 mins of imaging. Images show isolated mCherry from a single midplane section.
C. Time projection of mCherry signal over the 20 h 15 min duration of imaging. *Inset*, close-up of individual protrusions showing retraction (*top*) and extension (*bottom*).

D. Example midplane section showing overlay (*left*) and maximum intensity projection of GATA3 reporter (*middle*) from a z-stack (9 slices, 5 µm per slice) of a single iUB organoid acquired at day 12. Depth projection (*right*) shows the z-position of budding structures.
E. *Left*, manual ROI outline tracing. *Middle*, tip annotation for the example organoid shown in panel d. *Right*, the example organoid has 7 buds, a projected area of 147,545 $\mu m^2$, and circularity of 0.282. Data are representative of measurements used to generate **Fig. 1K-M**.
F. Schematic of the GDNF-RET perturbation experiment for iUB organoids. Organoids were grown under identical conditions from days 0-7 and embedded in Matrigel domes (see: **Methods**). From day 7-12, organoids in the negative control (-GDNF) condition received UBM with 0 ng $ml^{-1}$ GDNF. All other organoids received UBM with 50 ng $ml^{-1}$ from days 7-9. From days 9-12 we varied the concentration of GDNF and/or Selpercatinib (RETi). Organoids in the positive control group (+GDNF) received UBM with 50 ng $ml^{-1}$ GDNF, the excess GDNF group (++GDNF) received UBM with 250 ng $ml^{-1}$ GDNF, and the RET inhibitor group (+RETi) received UBM with 50 ng $ml^{-1}$ GDNF and 100 nM Selpercatinib.
G. Scatter plot of bud number (#) as a function of projected organoid area ($x10^4$ $\mu m^2$) for each condition in **Fig. 1K-M**. Data are pooled from 2 biological replicates, numbers (n) of organoids are given in figure panels.
H. All conditions from **Fig. S4G** plotted on a single axis. Data were fitted to a single linear regression model, $n_{tip}$ = 0.86(area) - 1.23.

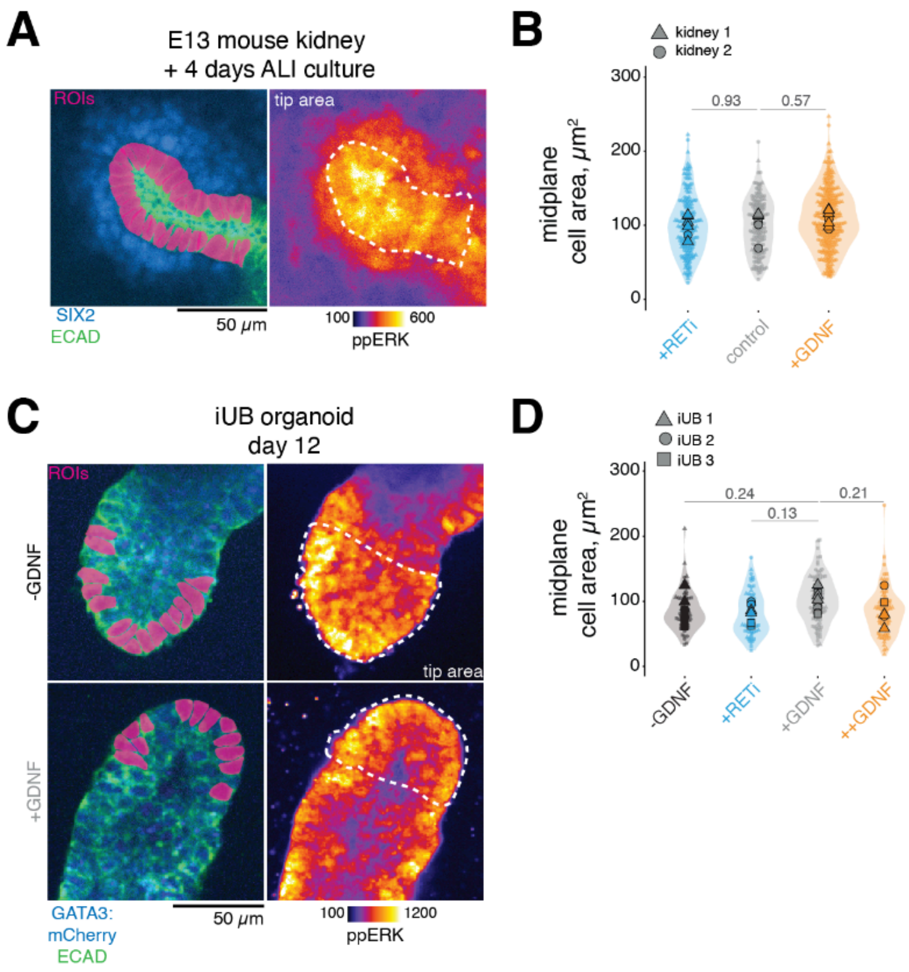


**Figure S5.** Immunofluorescence quantification of ERK in mouse and human tissues.

A. Example image of a ureteric bud tip in an E13 mouse explant after 4 days of ALI culture. *Left*, immunofluorescence shows locations of epithelial cell boundaries (ECAD) and the cap mesenchyme (SIX2), which defines the tip region. ROIs are manually drawn from individual epithelial cell boundaries. *Right*, intensity-coded ppERK (a.u.) shows activation across cap and tip cells. White dashed line shows the tip domain boundary/measurement area.

B. Midplane cell areas ($\mu m^2$) for ROIs in **Fig. 2B,C** for n = 197, 169, 312 tip cells measured from 7, 6, 8 tips (+RETi, control, +GDNF), N = 2 kidneys per condition. *P*-values by one-way ANOVA with Tukey's *post hoc* test, no significant comparisons ($p > 0.05$) between any conditions.

C. Example image of budding in day 12 iUB organoids in the -GDNF or +GDNF conditions (see also: **Fig. S4F** and **Table S1**). *Left*, immunofluorescence shows locations of epithelial cell boundaries (ECAD) and GATA3 reporter (mCherry). ROIs are manually drawn from individual epithelial cell boundaries. *Right*, intensity-coded ppERK (a.u.).

D. Midplane cell areas ($\mu m^2$) for ROIs in **Fig. 2E,F** for n = 67, 72, 97, 81 cells measured from 7, 9, 7, 6 tips (-GDNF, +RETi, +GDNF, ++GDNF). Data are collected from 3 organoids per condition. Dashed line shows microscope background cutoff (a.u.). *P*-values by one-way ANOVA with Dunnett's *post hoc* test using +GDNF as the reference group. No significant comparisons ($p > 0.05$) between any conditions.

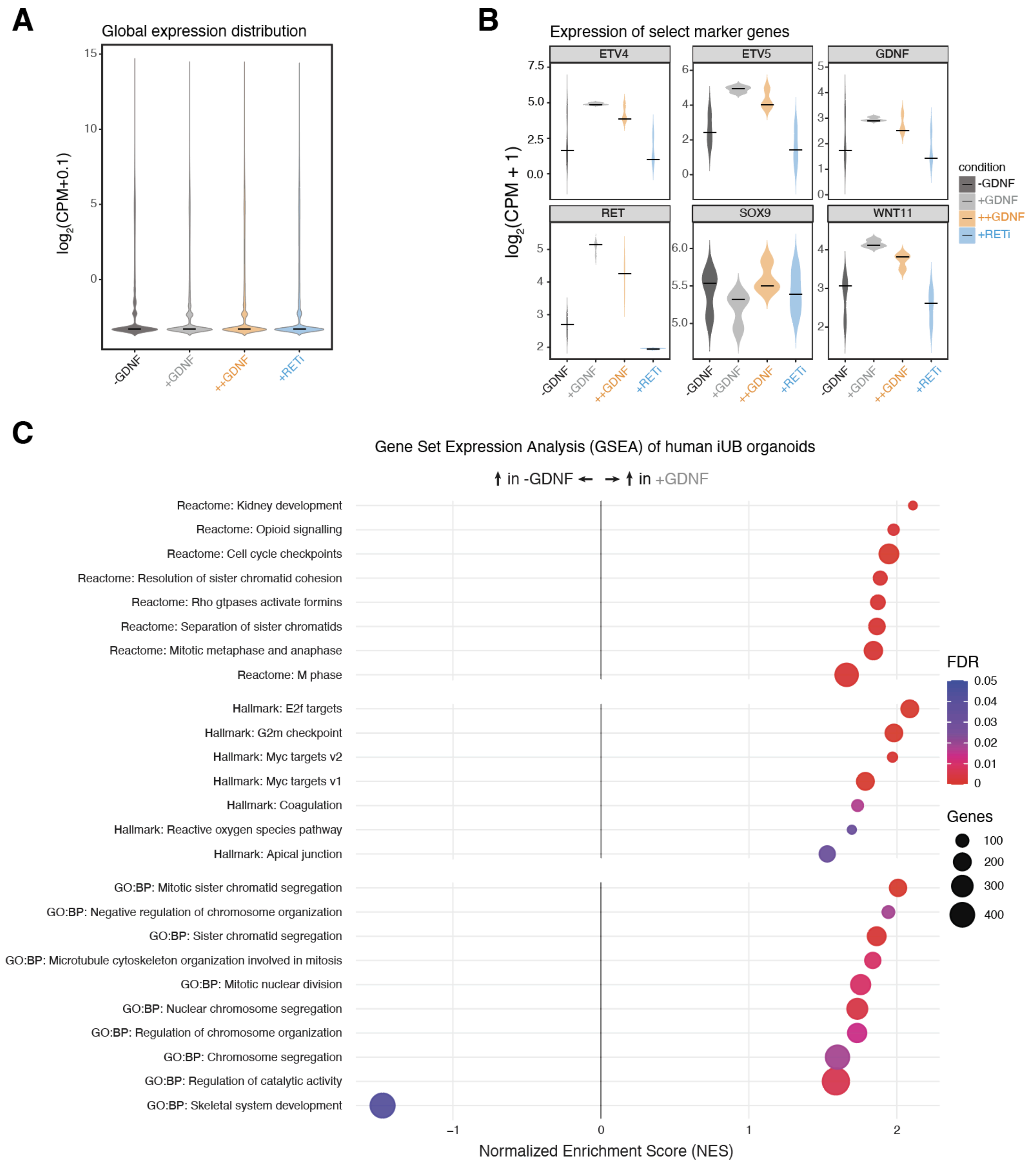


**Figure S6.** RNA-seq analysis of iUB organoids.

A. Global expression distribution of all genes in iUB organoids under the treatment conditions described in **Fig. 2I**. Gene expression is reported in $log_2$-transformed counts per million (CPM+0.1, a.u.).

B. Violin plots of ureteric bud tip markers (ETV4, ETV5, RET, WNT11, SOX9) and GDNF. Gene expression is reported in $log_2$-transformed counts per million (CPM+1, a.u.).

C. Gene Set Enrichment Analysis (GSEA) of upregulated genes in day 12 iUB organoids between the +GDNF and -GDNF groups as a function of normalized enrichment score (NES). Gene sets are color coded by false discovery rate (FDR), size coded by the size of the gene set. Data are derived from 3 biological replicates.

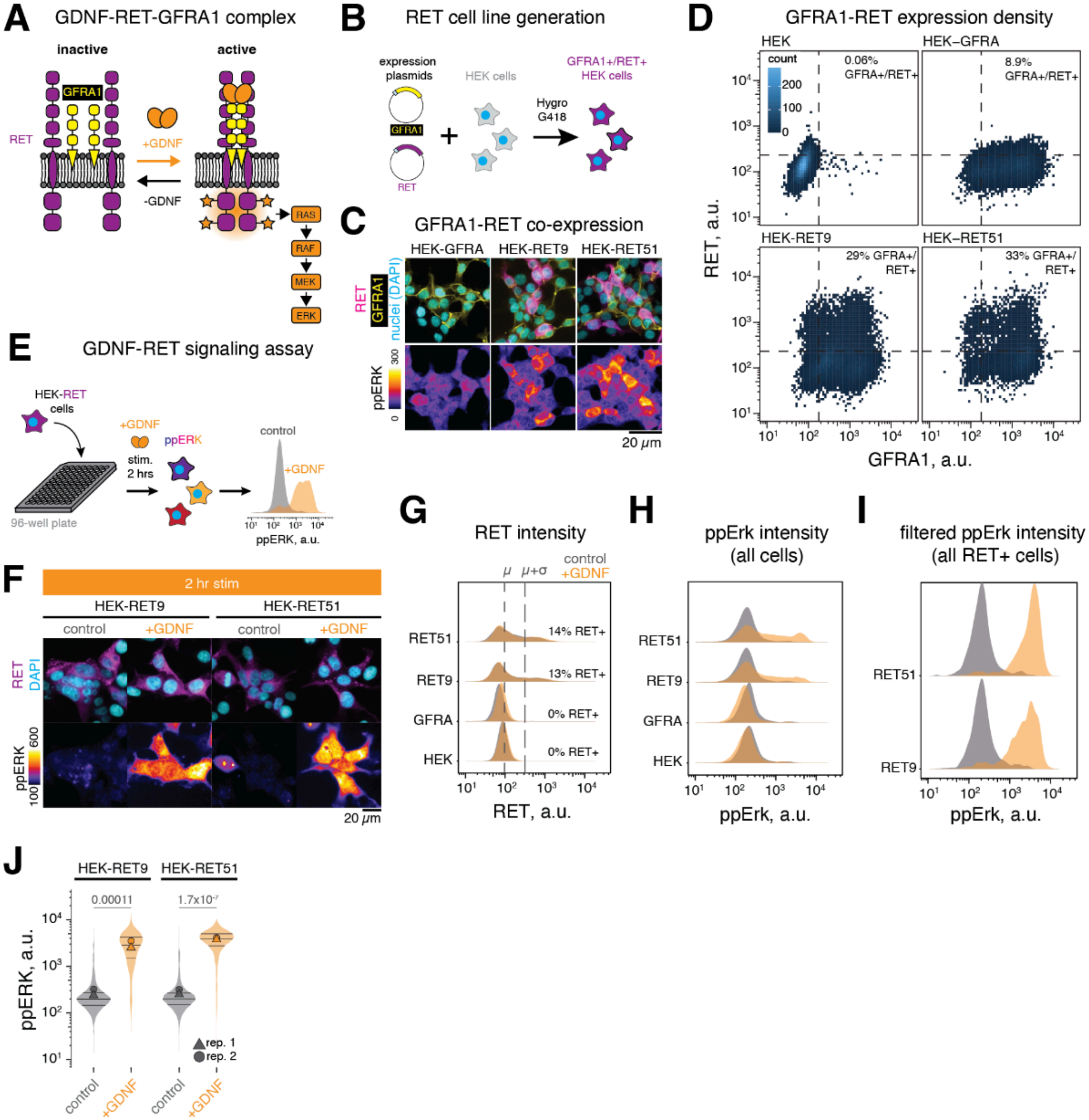


**Figure S7.** Quantitative ppERK analysis in HEK-RET cells.

A. Model for activation of the GDNF-RET-GFRA1 complex and downstream signaling through ERK.
B. Generation of HEK cell lines co-expressing GFRA1 and RET by transfection and antibiotic selection. HEK-GFRA cells express GFRA1 alone, HEK-RET cells co-express GFRA1 and either full-length RET9 or RET51.
C. Immunofluorescence of HEK-GFRA, HEK-RET9, and HEK-RET51 cells following 2 hr stimulation with 100 ng $ml^{-1}$ GDNF. Images in this panel were prepared using mouse anti-ppERK for multiplexing with other antibodies. Cells that co-express GFRA1 and RET have higher ppERK (a.u.) following stimulation.
D. Immunofluorescence density plot of RET and GFRA1 signal (a.u.) in HEK (parental line), HEK-GFRA1, HEK-RET9, and HEK-RET51 cells, n = 13186, 11101, 16618, 7227 cells (HEK, HEK-GFRA, HEK-RET9, HEK-RET51). Quadrants show mean + standard deviation ($\mu+\sigma$) cutoff

of signal in parental line, as well as percentages of RET+/GFRA1+ cells in each line (see: **Methods**).

E. GDNF stimulation of HEK-RET cells and density plot of ppERK (a.u.) from n = 1130, 1234 cells (control, +GDNF).
F. Immunofluorescence of HEK-RET9 and HEK-RET51 cells following 2 hr stimulation with 100 ng $ml^{-1}$ GDNF or control conditions. *Top*, RET and nuclei (DAPI). *Bottom*, intensity-coded ppERK (a.u.).
G. Mean ppERK intensity (a.u.) from RET+ cells across all conditions. HEK-RET9: n=1995, 2189 cells, HEK-RET51: n = 2483, 2913 (control, +GDNF). Data pooled from 2 biological replicates, each containing 3 technical replicates per condition. *P*-values by Welch's t-test. Images in this panel and **Fig. S8** were prepared using a rabbit anti-ppERK antibody.
H. Density plot of RET intensity (a.u.) for HEK (parental line), HEK-GFRA, HEK-RET9, and HEK-RET51 cells. Cells were serum-starved for 5-6 hours prior to stimulation. Vertical dashed lines represent the mean (μ) RET signal for all HEK parental cells and the RET+ intensity threshold, which is the mean plus one standard deviation of signal from HEK parental cells (μ+σ, see: **Methods**). HEK-RET9: n=1995, 2189 (control, +GDNF), HEK-RET51: n=2483, 2913 (control, +GDNF) cells, pooled from a single experiment containing 3 technical replicates.
I. Density plot of ppERK intensity (a.u.) for all cells above.
J. Density plot of ppERK intensity (a.u.) after filtering for RET+ cells, HEK-RET9: n = 1995, 2189 (control, +GDNF), HEK-RET51: n = 2483, 2913 (control, +GDNF) cells.

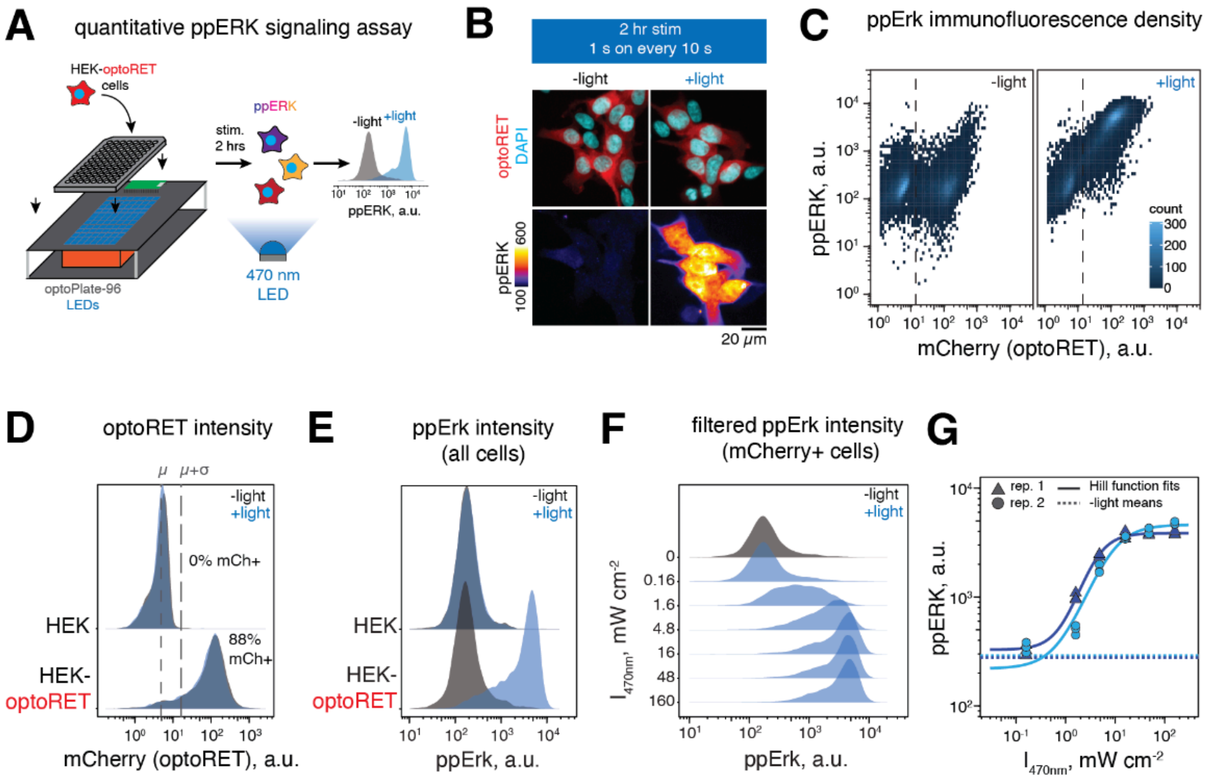


**H**

| $I_{470nm}$, mW cm$^{-2}$ | replicate 1 n (cells) | replicate 1 p-value vs. -light | replicate 2 n (cells) | replicate 2 p-value vs. -light |
|---|---|---|---|---|
| 0 (-light) | 4273 | na | 8274 | na |
| 0.16 | 8416 | 0.99 | 9630 | 0.99 |
| 1.6 | 6751 | 0.45 | 9842 | 0.0018 |
| 4.8 | 8139 | <0.0001 | 9560 | <0.0001 |
| 16 | 8036 | <0.0001 | 8232 | <0.0001 |
| 48 | 7364 | <0.0001 | 8535 | <0.0001 |
| 160 | 7427 | <0.0001 | 8499 | <0.0001 |

**Figure S8.** Quantitative ppERK analysis in HEK-optoRET cells.

A. Blue light stimulation of HEK-optoRET cells in the optoPlate-96 and density plot of ppERK (a.u.) from n = 4447, 7759 cells (dark, +light).
B. Immunofluorescence of HEK-optoRET cells following 2 hr stimulation with 470 nm light at 0 mW cm$^{-2}$ (-light) or 150 mW cm$^{-2}$ for 1 s every 10 s (+light). *Top*, optoRET(mCherry) and nuclei (DAPI). *Bottom*, intensity-coded ppERK (a.u.).
C. Immunofluorescence density plot of ppERK (a.u.) as a function of optoRET(mCherry) intensity (a.u.) in HEK and HEK-optoRET cells stimulated with 470 nm light at 160 mW cm$^{-2}$ (+light) or 0 mW cm$^{-2}$ (-light) for 1 s every 10 s for 2 hrs, n = 13975, 17920 cells (-light, +light) from 2 biological replicates, each containing 3 technical replicates
D. Density plot of optoRET (mCherry) intensity (a.u.) in HEK (parental line) and HEK-optoRET cells stimulated with 470 nm light at 160 mW cm$^{-2}$ (+light) or 0 mW cm$^{-2}$ (-light) for 1 s every 10 s for 2 hrs using an optoPlate-96 [52]. Cells were starved for 6 hours prior to stimulation Vertical dashed lines represent the mean (μ) of all HEK parental cells and the optoRET+ intensity threshold (μ+σ of all untransfected cells). HEK: n = 10832, 5752 (-light, +light), HEK-optoRET: n = 13975, 17920 (-light, +light) cells, pooled from 2 biological replicates each containing 3 technical replicates.

E. Density plot of ppERK (a.u.) for all cells above before filtering by optoRET(mCherry) expression.
F. Density plot of ppERK (a.u.) for all filtered optoRET+ cells above. After filtering by optoRET(mCherry) expression, n = 4273, 8416, 6751, 8139, 8036, 7364, 7427 cells (0, 0.16, 0.48, 1.6, 4.8, 16, 48, 160 mW $cm^{-2}$) from a single experiment containing 3 technical replicates per condition.
G. Violin plot of ppERK (a.u.) for all filtered optoRET+ cells above.
H. Numbers (n) of cells and statistical tests for all HEK-optoRET cells above. *P*-values by one-way ANOVA with Dunnett's *post hoc* test using -light (0 mW $cm^{-2}$) as the reference condition.

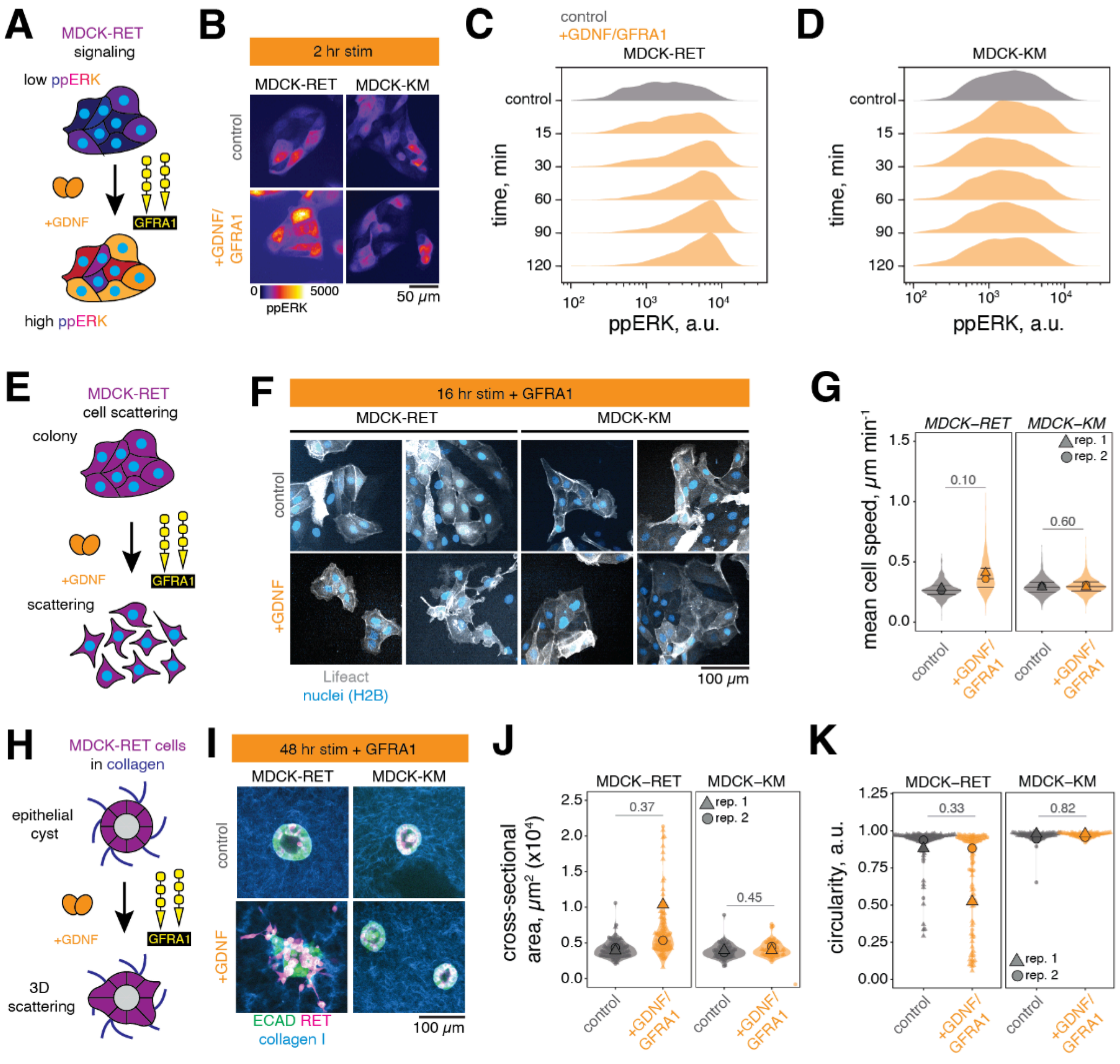


**Figure S9.** Quantification of ERK signaling, scattering, and morphogenesis in MDCK-RET and MDCK-KM cells.

A. MDCK-RET signaling response to GDNF+GFRA1.
B. Immunofluorescence images of nuclei (DAPI) and intensity-coded ppERK (a.u.) in MDCK-RET or MDCK-KM cells. MDCK-KM cells express a full-length RET construct with a point mutation (K758M) that renders the tyrosine kinase domain nonfunctional and are suitable as negative controls [54]. Cells were treated with 50 ng ml$^{-1}$ GDNF and 100 ng ml$^{-1}$ GFRA1 or left untreated (control) for 2 hours before fixing and immunostaining.
C. Density plot of ppERK (a.u.) in MDCK-RET cells under control conditions or after defined stimulation intervals (15’, 30’, 60’, 90’, 120’) with GDNF/GFRA1, n = 4468, 4108, 4528, 4347, 4510, 4411 cells (control, 15’, 30’, 60’, 90’, 120’). Data are pooled from 3 biological replicates each containing 3 technical replicates each.
D. Density plot of ppERK (a.u.) in MDCK-KM cells under control conditions or after defined stimulation intervals (15’, 30’, 60’, 90’, 120’) with GDNF/GFRA1, n = 3538, 4604, 4452, 4952, 4619, 4911 cells (control, 15’, 30’, 60’, 90’, 120’). Data are pooled from 3 biological replicates

each containing 3 technical replicates each.

E. MDCK-RET scattering response to GDNF+GFRA1.

F. Time lapse image sequences of MDCK-RET or MDCK-KM cells co-expressing Lifeact-eGFP and H2B-mRuby2. Cells were treated with 50 ng $ml^{-1}$ GDNF or left untreated (control) and imaged every 5 min for 12-16 hours. All conditions received 100 ng $ml^{-1}$ GFRA1.

G. Mean cell speed for single MDCK-RET or MDCK-KM cells under control or +GDNF conditions. MDCK-RET: n = 435, 524 cells (control, +GDNF); MDCK-KM: n = 1151, 1201 cells (control, +GDNF). Data are pooled from 2 biological replicates each containing 2 technical replicates. *P*-values by Welch's t-test.

H. MDCK-RET cyst morphogenesis response to GDNF+GFRA1.

I. Immunofluorescence images of MDCK-RET and MCDK-KM cysts following stimulation for 48 hours with 50 ng $ml^{-1}$ GDNF or left untreated (control) showing ECAD and RET expression. Collagen I fibers were pre-labeled with NHS-Alexa 647. All conditions received 100 ng $ml^{-1}$ GFRA1.

J. Cross-sectional area ($x10^4$ $\mu m^2$) for MDCK-RET and MDCK-KM cysts under control or +GDNF conditions. MDCK-RET: n = 109, 111 cysts (control, +GDNF); MDCK-KM: n = 76, 68 cysts (control, +GDNF). Data are pooled from 2 biological replicates.

K. Circularity (a.u.) for MDCK-RET and MDCK-KM cysts. *P*-values in panels j, k by Welch's t-test.

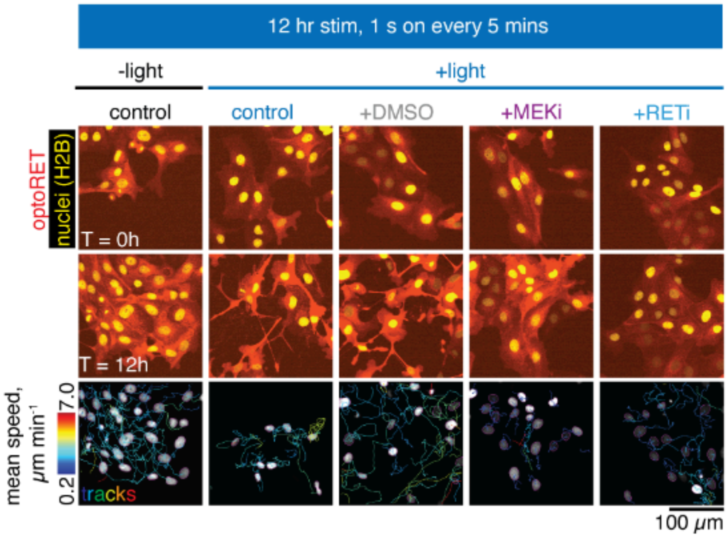


**Figure S10.** MDCK-optoRET scattering under different signaling conditions.

Images of MDCK-optoRET cells co-expressing H2B-mVenus. *Top row*, T = 0 h. *Middle*, T = 12 hrs stimulation with 488 nm light (2 mW, 1 s every 5 mins) or left unstimulated. Stimulated cells also received concurrent treatment with DMSO (vehicle), 100 nM Trametinib (+MEKi), or 100 nM Selpercatinib (+RETi) for the duration of imaging. *Bottom*, TrackMate [56] outputs acquired fromH2B stacks for all conditions above using the StarDist algorithm [85]. Tracks are color-coded by cell speed.

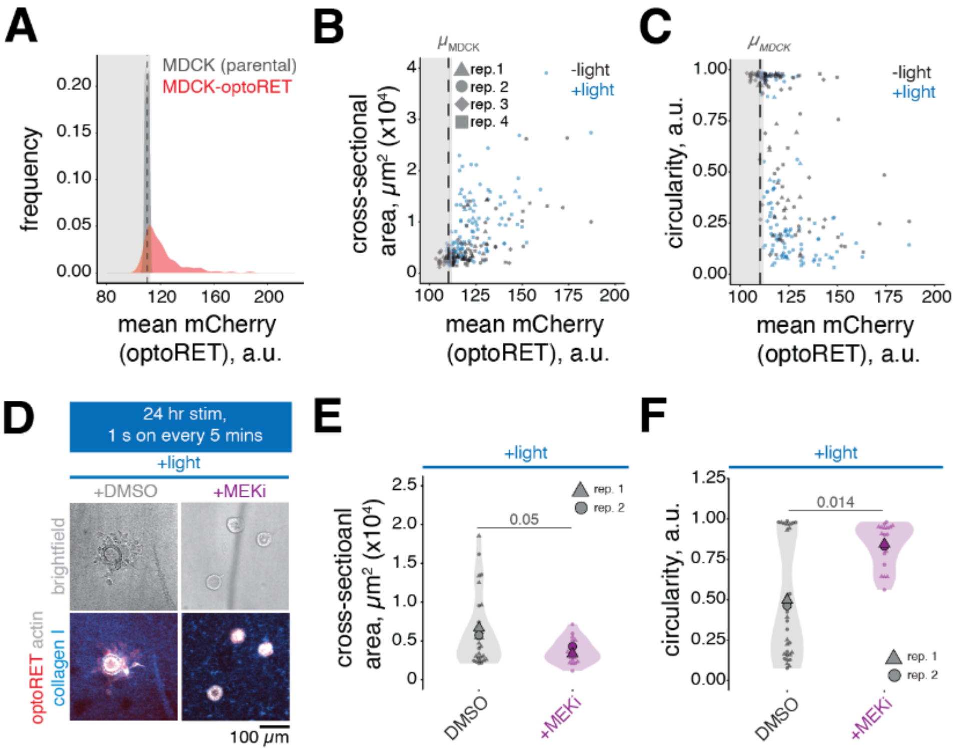


**Figure S11.** Quantitative metrics of MDCK cyst morphogenesis.

A. Density plot of mean mCherry expression (a.u.) for all MDCK and MDCK-optoRET cysts in the experiment in **Fig. 3K-M**, n = 184 MDCK and 266 MDCK-optoRET cysts pooled from 4 biological replicates.
B. Scatter plot of cross sectional area (μm$^2$ x10$^4$) as a function of mean mCherry expression (a.u.) for all MDCK-optoRET cysts in **Fig. 3K-M**. Vertical dashed line indicates a mean plus standard deviation (μ+σ) minimum cutoff for mean mCherry expression measured from MDCK cells (see: **Methods**).
C. Scatter plot of circularity (a.u.) as a function of mean mCherry expression (a.u.) for MDCK-optoRET cysts.
D. Immunofluorescence images of MDCK-optoRET cysts following 24 hr stimulation with 470 nm light (50 mW cm$^{-2}$, 1 s every 5 mins) in an optoPlate-96. *Top*, brightfield. *Bottom*, optoRET (mCherry), actin (phalloidin), and labeled collagen I. Cysts were treated with DMSO or 100 nM Trametinib (+MEKi) for the duration of stimulation.
E. Cross-sectional area (μm$^2$ x10$^4$) for MDCK-optoRET cysts, n = 30, 20 (DMSO, +MEKi) cysts pooled from 2 biological replicates. Data were filtered for mean mCherry expression (a.u.) using MDCK controls (see: **Methods**).
F. Circularity (a.u.) for all MDCK-optoRET cysts treated with DMSO or +MEKi. *P*-values in panels e, f by Wilcoxon rank-sum test.

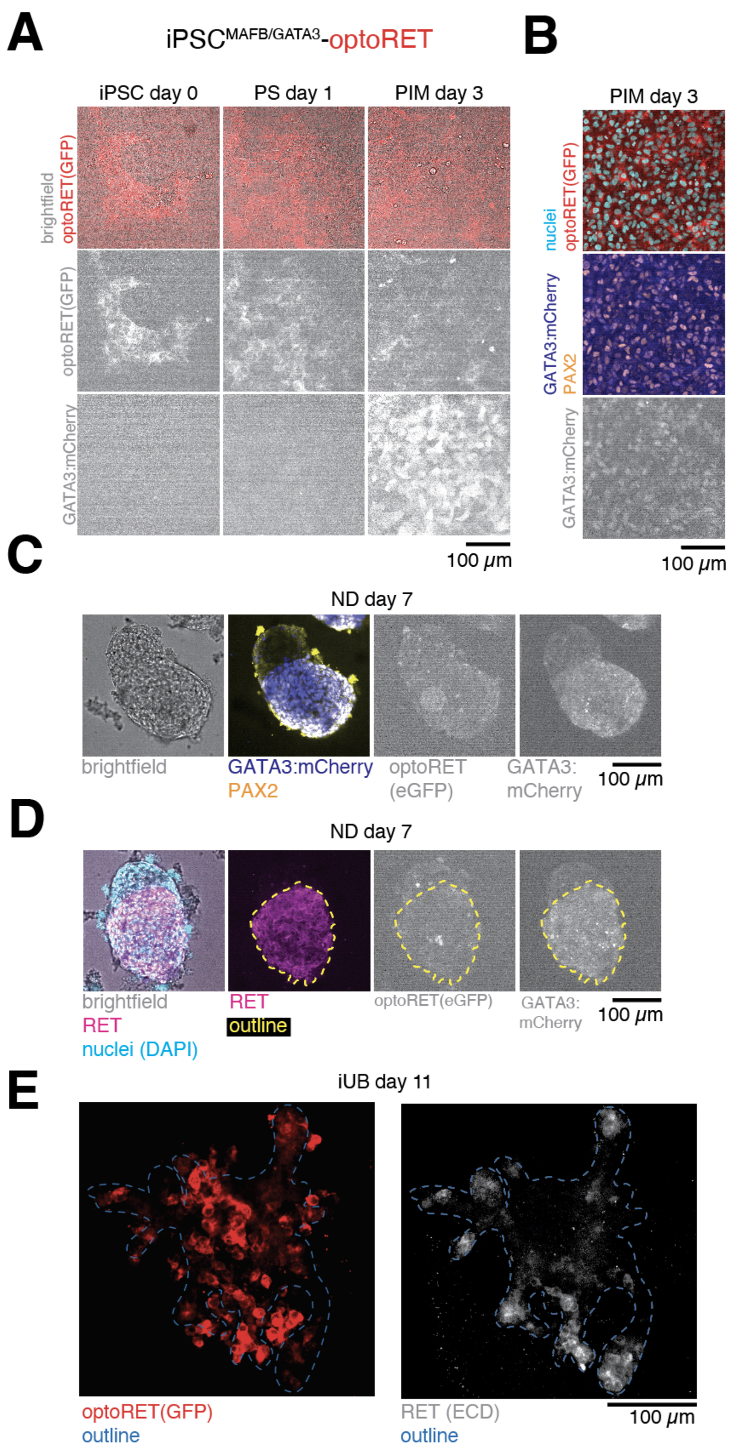


**Figure S12.** Differentiation of iPSC-optoRET cells into iUB organoids and spatial disposition of activated optoRET+ cells.

A. Cell lines stably expressing an eGFP-tagged optoRET were generated from iPSC$^{MAFB/GATA3}$ cells (*left column*, iPSC-optoRET) using PiggyBac transposase, purified by FACS, and differentiated into day 3 PIM cells (*middle* and *right columns*, **Fig. 4** and **Methods**). *Top row*, brightfield and optoRET (mCherry) overlay. *Middle row*, isolated optoRET(eGFP). *Bottom row*, isolated GATA3 reporter (mCherry) expression increases over days 0-3 indicating differentiation towards the PIM lineage.

B. Immunofluorescence of day 3 PIM-optoRET cells shows robust co-expression of PAX2, and GATA3:mCherry in PIM-optoRET cells. *Top*, optoRET and nuclei (DAPI). *Middle*, GATA3 reporter (mCherry) and PAX2 immunofluorescence. *Bottom*, isolated GATA3 reporter (mCherry).
C. Immunofluorescence of day 7 optoRET+ iUB/ND spheroids shows continued co-expression of PAX2 and GATA3 and sorting into a GATA3+ epithelium and GATA3- stromal population.
D. Immunofluorescence of day 7 optoRET+ iUB/ND spheroids shows robust expression of full-length RET within the GATA3+ epithelial population. Immunostaining was performed using a commercially available goat anti-human RET antibody raised against the extracellular domain (aa29-615), which are not present in the optoRET(eGFP) construct.
E. Immunofluorescence of day 11 optoRET(GFP) and RET(ECD) after 4 days of DMD stimulation (+light whole, **Fig. 5**).

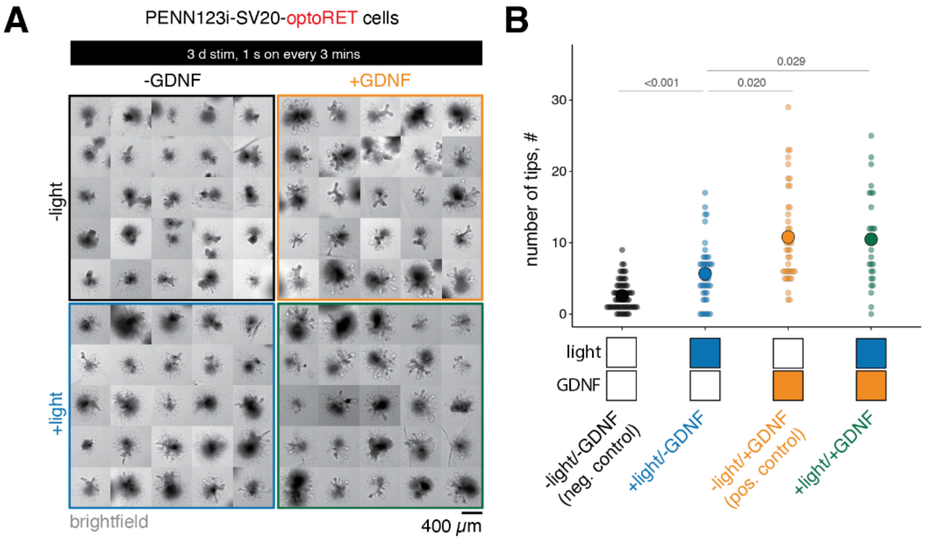


**Figure S13.** Optogenetic budding in SV20-derived iUB organoids.

A. Representative gallery of day 10 iUB-optoRET organoids derived from SV20 iPSCs illustrating morphology outcomes across −light/+light and −GDNF/+GDNF conditions. Cells were cultured from days 0-7 and plated in Matrigel droplets in a 6-well plate as described in **Methods**. Droplets were aligned to arrays of 470 nm LEDs on an optoPlate-96 and organoids were stimulated for 1 s every 3 mins at 50 mW $cm^{-2}$ over 3 days for the +light conditions. Organoids in the -light/-GDNF (control) and -light/+GDNF conditions were cultured in a separate plate to prevent light exposure. All plates were wrapped in aluminum foil for the duration of the experiment to prevent outside light exposure.

B. Bud number on day 10 across all conditions, n = 65, 39, 40, 28 organoids (control, +light, +GDNF, +light/+GDNF) from a single experiment. *P*-values by one-way Kruskal-Wallis test with Dunn's *post hoc* test.

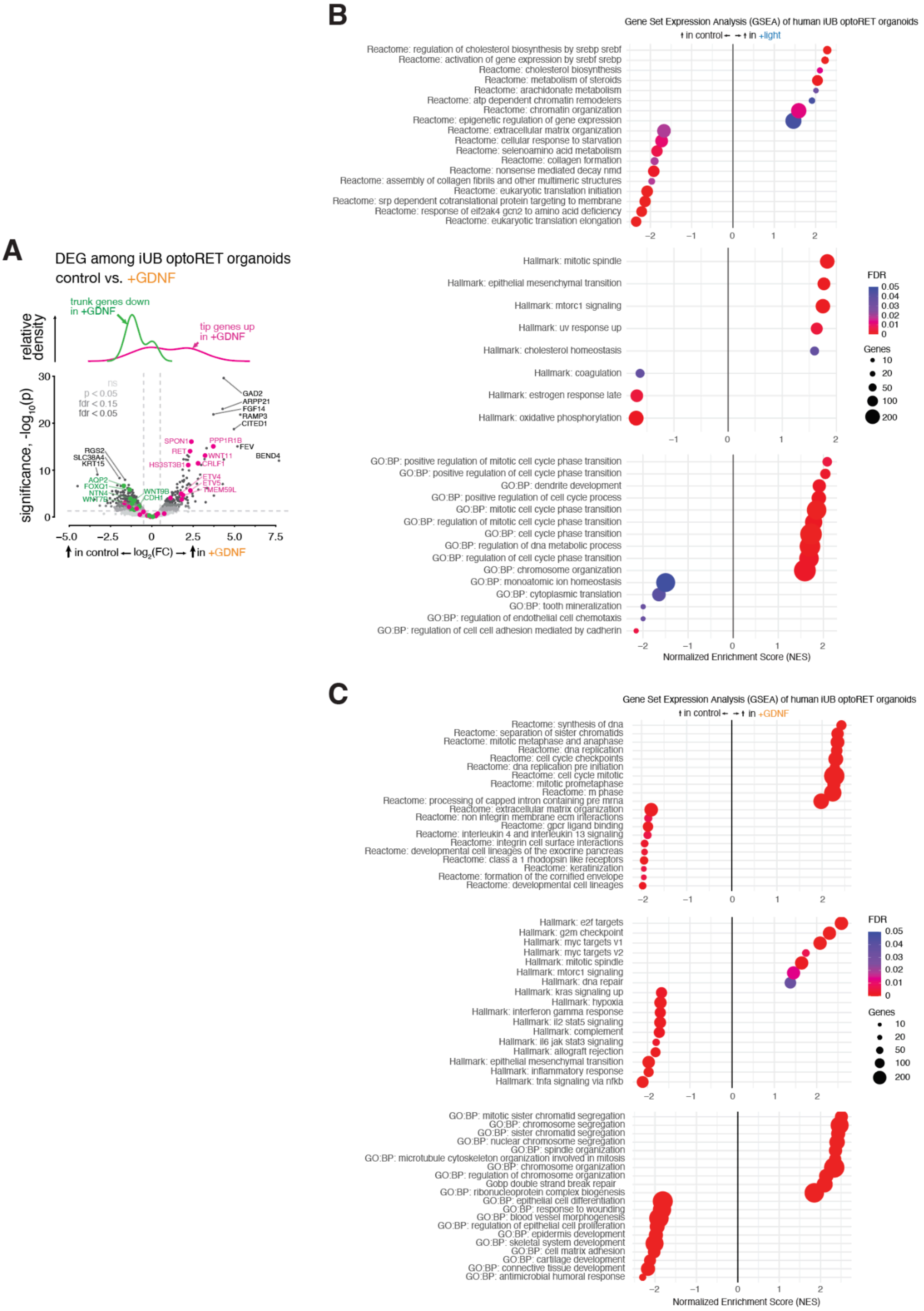
A
DEG among iUB optoRET organoids
control vs. +GDNF
trunk genes down
in +GDNF
tip genes up
in +GDNF
relative density
significance, -log10(p)
ns
p < 0.05
fdr < 0.15
fdr < 0.05
GAD2
ARPP21
FGF14
RAMP3
CITED1
PPP1R1B
FEV
BEND4
SPON1
RET
WNT11
HS3ST3B1
CRLF1
ETV4
ETV5
TMEM59L
RGS2
SLC38A4
KRT15
AQP2
FOXQ1
NTN4
WNT7B
WNT9B
CDH1
in control ← log2(FC) → in +GDNF
B
Gene Set Expression Analysis (GSEA) of human iUB optoRET organoids
in control ← → in +light
Reactome: regulation of cholesterol biosynthesis by srebp srebf
Reactome: activation of gene expression by srebf srebp
Reactome: cholesterol biosynthesis
Reactome: metabolism of steroids
Reactome: arachidonate metabolism
Reactome: atp dependent chromatin remodelers
Reactome: chromatin organization
Reactome: epigenetic regulation of gene expression
Reactome: extracellular matrix organization
Reactome: cellular response to starvation
Reactome: selenoamino acid metabolism
Reactome: collagen formation
Reactome: nonsense mediated decay nmd
Reactome: assembly of collagen fibrils and other multimeric structures
Reactome: eukaryotic translation initiation
Reactome: srp dependent cotranslational protein targeting to membrane
Reactome: response of eif2ak4 gcn2 to amino acid deficiency
Reactome: eukaryotic translation elongation
Hallmark: mitotic spindle
Hallmark: epithelial mesenchymal transition
Hallmark: mtorc1 signaling
Hallmark: uv response up
Hallmark: cholesterol homeostasis
Hallmark: coagulation
Hallmark: estrogen response late
Hallmark: oxidative phosphorylation
FDR
Genes
GO:BP: positive regulation of mitotic cell cycle phase transition
GO:BP: positive regulation of cell cycle phase transition
GO:BP: dendrite development
GO:BP: positive regulation of cell cycle process
GO:BP: mitotic cell cycle phase transition
GO:BP: regulation of mitotic cell cycle phase transition
GO:BP: cell cycle phase transition
GO:BP: regulation of dna metabolic process
GO:BP: regulation of cell cycle phase transition
GO:BP: chromosome organization
GO:BP: monoatomic ion homeostasis
GO:BP: cytoplasmic translation
GO:BP: tooth mineralization
GO:BP: regulation of endothelial cell chemotaxis
GO:BP: regulation of cell cell adhesion mediated by cadherin
Normalized Enrichment Score (NES)
C
Gene Set Expression Analysis (GSEA) of human iUB optoRET organoids
in control ← → in +GDNF
Reactome: synthesis of dna
Reactome: separation of sister chromatids
Reactome: mitotic metaphase and anaphase
Reactome: dna replication
Reactome: cell cycle checkpoints
Reactome: dna replication pre initiation
Reactome: cell cycle mitotic
Reactome: mitotic prometaphase
Reactome: m phase
Reactome: processing of capped intron containing pre mrna
Reactome: extracellular matrix organization
Reactome: non integrin membrane ecm interactions
Reactome: gpcr ligand binding
Reactome: interleukin 4 and interleukin 13 signaling
Reactome: integrin cell surface interactions
Reactome: developmental cell lineages of the exocrine pancreas
Reactome: class a 1 rhodopsin like receptors
Reactome: keratinization
Reactome: formation of the cornified envelope
Reactome: developmental cell lineages
Hallmark: e2f targets
Hallmark: g2m checkpoint
Hallmark: myc targets v1
Hallmark: myc targets v2
Hallmark: mitotic spindle
Hallmark: mtorc1 signaling
Hallmark: dna repair
Hallmark: kras signaling up
Hallmark: hypoxia
Hallmark: interferon gamma response
Hallmark: il2 stat5 signaling
Hallmark: complement
Hallmark: il6 jak stat3 signaling
Hallmark: allograft rejection
Hallmark: epithelial mesenchymal transition
Hallmark: inflammatory response
Hallmark: tnfa signaling via nfkb
FDR
Genes
GO:BP: mitotic sister chromatid segregation
GO:BP: chromosome segregation
GO:BP: sister chromatid segregation
GO:BP: nuclear chromosome segregation
GO:BP: spindle organization
GO:BP: microtubule cytoskeleton organization involved in mitosis
GO:BP: chromosome organization
GO:BP: regulation of chromosome organization
Gobp double strand break repair
GO:BP: ribonucleoprotein complex biogenesis
GO:BP: epithelial cell differentiation
GO:BP: response to wounding
GO:BP: blood vessel morphogenesis
GO:BP: regulation of epithelial cell proliferation
GO:BP: epidermis development
GO:BP: skeletal system development
GO:BP: cell matrix adhesion
GO:BP: cartilage development
GO:BP: connective tissue development
GO:BP: antimicrobial humoral response
Normalized Enrichment Score (NES)

**Figure S14.** Supplementary analysis for optoRET organoid bulk RNA-seq.

A. Differential expression analysis from bulk RNA-seq of iUB-optoRET organoids grown in ±GDNF stimulation conditions. *Top*, relative density of tip and trunk marker genes. *Bottom*, volcano plot of all genes organized by significance (-log10(p)) and log2(fold-change) with tip and trunk-specific markers highlighted. Data are derived from 3 biological replicates.
B. Full GSEA analysis for ±light stimulation conditions.
C. Full GSEA analysis for ±GDNF stimulation conditions.

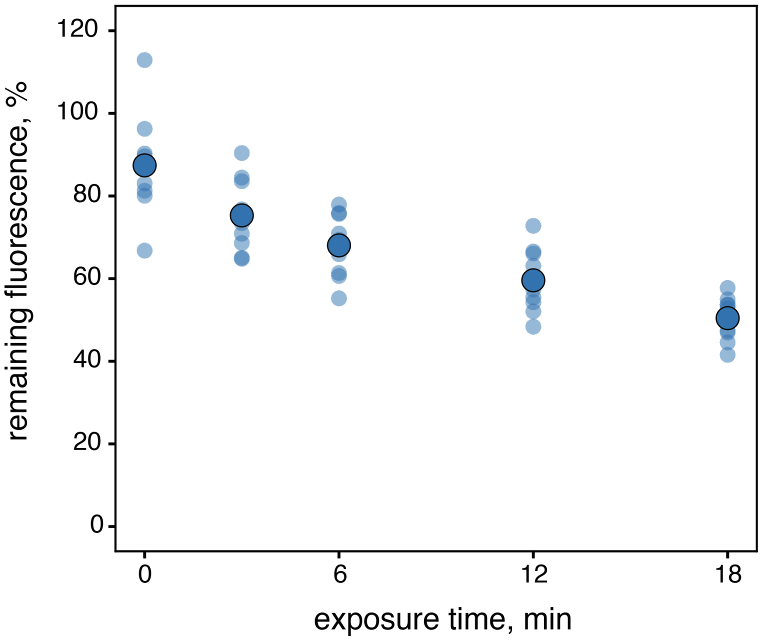


**Figure S15.** Calibration of DMD illumination intensity by fluorescent bead photobleaching. Calibration curve of 10 µm FITC-labeled polystyrene microspheres showing % FITC fluorescence remaining after the indicated continuous exposure time at 5% DMD LED power. By comparison, >90% fluorescence remained for beads similarly exposed in the optoPlate-96 at ~100% power (320 mW $cm^{-2}$ nominal power) after 12 min. This indicates that optoPlate-96 exposure is sufficient to induce morphogenesis due to optoRET activation, while falling below a power suitable for direct calibration against the DMD.

### Supplementary Tables

#### **Table S1.** Differentiation media for iUB

| Stage | Day | Basal Medium | Culture Forn | Y-27632 | Activin A | BMP4 | CHIR | FGF2 | A83-01 | LDN | RA | FGF9 | GDNF | FGF10 | Selpercatinib |
|---|---|---|---|---|---|---|---|---|---|---|---|---|---|---|---|
| iPSC seeding | -1 | mTeSR+ | Monolayer | 10μM | - | - | - | - | - | - | - | - | - | - | - |
| Primitive streak | 0-1 | Adv. RPMI + Glu | Monolayer | - | 50ng $mL^{-1}$ | 25ng $mL^{-1}$ | 5μM | 25ng $mL^{-1}$ | - | - | - | - | - | - | - |
| Pronephric IM | 1-3 | Adv. RPMI + Glu | Monolayer | - | - | - | - | 25ng $mL^{-1}$ | 1μM | 0.1μM | 0.1μM | - | - | - | - |
| ND spheroid | 3-5 | Adv. RPMI + Glu | Aggrewell | - | - | - | - | - | - | - | 0.1μM | 50ng $mL^{-1}$ | - | - | - |
| ND spheroid | 5-7 | Adv. RPMI + Glu | Aggrewell | - | - | - | - | - | - | - | 0.1μM | - | 50ng $mL^{-1}$ | - | - |
| UBM (Standard) | 7-12 | Adv. RPMI + Glu | Matrigel dropl | 10μM | - | - | 2uM | - | 1μM | 0.1μM | 0.1μM | - | 50ng $mL^{-1}$ | 50ng $mL^{-1}$ | - |
| UBM (-GDNF) | 7-12 | Adv. RPMI + Glu | Matrigel dropl | 10μM | - | - | 2uM | - | 1μM | 0.1μM | 0.1μM | - | - | 50ng $mL^{-1}$ | - |
| UBM (++GDNF) | 9-12 | Adv. RPMI + Glu | Matrigel dropl | 10μM | - | - | 2uM | - | 1μM | 0.1μM | 0.1μM | - | 250ng $mL^{-1}$ | 50ng $mL^{-1}$ | - |
| UBM (+RETi) | 9-12 | Adv. RPMI + Glu | Matrigel dropl | 10μM | - | - | 2uM | - | 1μM | 0.1μM | 0.1μM | - | 50ng $mL^{-1}$ | 50ng $mL^{-1}$ | 0.1μM |

#### **Table S2.** Primary and secondary antibodies

| Antibody | Species | Dilution | Manufacturer | Cat# | RRID |
|---|---|---|---|---|---|
| E-cadherin (36/E) | Mouse | 1:500 (organoids) | BD Biosciences | #610182 | AB_397581 |
| E-cadherin (24E10) | Rabbit | 1:200 | Cell Signaling Technologies | #3195 | AB_2291471 |
| E-cadherin (DECMA-1) | Rat | 1:200 | Abcam | #ab11512 | AB_1210458 |
| EGFP | Chicken | 1:2000 | Abcam | #ab13970 | AB_300798 |
| GATA-3 | Goat | 1:200 | R&D Systems | #AF2605 | AB_2108571 |
| GFR alpha-1/GDNF R alpha-1 | Goat | 1:200 | R&D Systems | #AF714 | AB_355541 |
| Integrin alpha 8 (ITGA8) | Goat | 1:300 (explants) | R&D Systems | #AF4076 | AB_2296280 |
| Jagged 1 | Goat | 1:200 (explants) | R&D Systems | #AF599 | AB_2128257 |
| Cytokeratin-8 (C51) | Mouse | 1:200 | Santa Cruz | #sc-8020 | AB_627857 |
| Nanog [23D2-3C6] | Mouse | 1:200 | Abcam | #ab173368 | AB_3076592 |
| PAX2 | Rabbit | 1:200 | Invitrogen | #71-6000 | AB_2533990 |
| Phospho-p44/42 MAPK (ppErk 1/2) | Mouse | 1:400 | Cell Signaling Technologies | #5726 | AB_2797617 |
| Phospho-p44/42 MAPK (ppErk 1/2) | Rabbit | 1:400 (cells), 1:200 (explants/organoids) | Cell Signaling Technologies | #4370 | AB_2315112 |
| RET (C31B4) | Rabbit | 1:200 | Cell Signaling Technologies | #3223 | AB_2238465 |
| hRET | Goat | 1:400 (cells), 1:200 (organoids) | R&D Systems | #AF1485 | AB_354820 |
| mRET | Goat | 1:100 (explants) | R&D Systems | #AF482 | AB_2301030 |
| SIX2 (monoclonal) | Mouse | 1:250 (explants) | Proteintech | #66347-1-Ig | AB_2881727 |
| SIX2 (polyclonal) | Rabbit | 1:600 (explants) | Proteintech | #11562–1-AP | AB_2189084 |
| TBXT/Brachyury | Goat | 1:200 | R&D Systems | #AF2085 | AB_2200235 |
| Anti-chicken AlexaFluor™ 488 | Donkey | see methods | ThermoFisher | #A78948 | AB_2921070 |
| Anti-goat AlexaFluor™ 555 | Donkey | see methods | ThermoFisher | #A21432 | AB_2535853 |
| Anti-goat AlexaFluor™ Plus 647 | Donkey | see methods | ThermoFisher | #A32849 | AB_276840 |
| Anti-mouse AlexaFluor™ 488 | Donkey | see methods | ThermoFisher | #A21202 | AB_141607 |
| Anti-mouse AlexaFluor™ Plus 647 | Donkey | see methods | ThermoFisher | #A31571 | AB_162542 |
| Anti-rabbit AlexaFluor™ 488 | Donkey | see methods | ThermoFisher | #A21206 | AB_2535792 |
| Anti-rabbit AlexaFluor™ 555 | Donkey | see methods | ThermoFisher | #A32794 | AB_2762834 |
| Anti-rat AlexaFluor™ Plus 405 | Donkey | see methods | ThermoFisher | #A48268 | AB_2890549 |

### Supplementary Movies

**Movie S1. Budding and branching of Day 10 iUB organoids in response to GDNF.** Representative confocal fluorescence timelapses of organoids in -GDNF (control) and +GDNF conditions. Images stacks were acquired at 10x magnification at 45 minute intervals over 20 hrs 15 mins. GATA3:mCherry reporter, blue; Brightfield, gray.

**Movie S2. Budding and branching of Day 9 iUB organoids in response to GDNF.** Representative epifluorescence timelapse of organoids in +GDNF condition acquired at 10x magnification. Image stacks were acquired at 10x magnification at 2 hr intervals over 54 hrs. GATA3:mCherry reporter, blue; Brightfield, gray.

**Movie S3. Scattering of MDCK-RET and MDCK-KM cells in response to GDNF.** Representative confocal fluorescence timelapses of MDCK-RET and MDCK-KM scattering in control and +GDNF conditions. Frames were acquired at 20x magnification at 5 min intervals over 16 hrs. All conditions received 100 ng $ml^{-1}$ Gfrα1 for the duration of imaging. H2B-mRuby2, blue; Lifeact-eGFP, gray.

**Movie S4. Scattering of MDCK-optoRET cells in response to light - 12 hr timelapse.** Representative confocal fluorescence fields of view for MDCK-optoRET cell scattering in the indicated treatment groups. Frames were acquired at 20x magnification at 5 min intervals over 12 hrs with a 488 nm laser pulse (20% power, 1 s) following each frame. MDCK-optoRET, red; H2B-mVenus, yellow.

**Movie S5. Symmetry breaking of 3D MDCK-RET cysts in response to GDNF.** Representative confocal fluorescence timelapse of MDCK-RET cysts in collagen I in the control and +GDNF conditions. Frames were acquired at 20x magnification at 30 min intervals over 24 hrs. All conditions received 100 ng $ml^{-1}$ Gfrα1 for the duration of imaging. Collagen I-AF647, magenta; H2B-mRuby2, blue; Lifeact-eGFP, gray.

**Movie S6. Symmetry breaking of 3D MDCK-optoRET cyst in response to light.** Confocal fluorescence timelapse of an MDCK-optoRET cyst in collagen I in the +light condition. Frames were acquired at 20x magnification at 30 min intervals over 24 hrs with a 488 nm laser stimulation loop (20% power, 1 s every 5 mins) run after each timepoint. Collagen I-AF647, magenta; Lifeact-mCherry, gray.

**Movie S7. OptoRET clustering in SV20 iPSCs in response to light.** Representative confocal fluorescence timelapse of Myr-RET(ICD)-eGFP-$CRY2^{PHR}$ (gray) clustering in SV20 iPSCs during blue light exposure. Frames were acquired at 60x magnification at 1 min intervals over 10 mins.

**Movie S8: Budding and branching of Day 7 optoRET-iUB organoids in response to light delivered by DMD.** Representative confocal fluorescence timelapses for organoids in +light whole (*left*) vs. +light half (*right*) DMD exposure conditions. Image stacks were acquired at 20x magnification at 2 hr intervals over 60 hrs with a 488 nm DMD stimulation loop (0.5 s every 4 min) run after each timepoint. Stage *xyz* positions were periodically updated to align organoids within the fixed ROI (cyan). GATA3:mCherry reporter, blue; Brightfield, gray.
